# Synthetic microbial sensing and biosynthesis of amaryllidaceae alkaloids

**DOI:** 10.1101/2023.04.05.535710

**Authors:** Simon d’Oelsnitz, Daniel Diaz, Daniel Acosta, Mason Schechter, Matthew Minus, James Howard, James Loy, Hannah Do, Hal S. Alper, Andrew D. Ellington

**Author notes:** To whom correspondence should be addressed: Simon d’Oelsnitz.

## Abstract

A major challenge to achieving industry-scale biomanufacturing of therapeutic alkaloids is the slow process of biocatalyst engineering. Amaryllidaceae alkaloids, such as the Alzheimer’s medication galantamine, are complex plant secondary metabolites with recognized therapeutic value. Due to their difficult synthesis they are regularly sourced by extraction and purification from low-yielding plants, including the wild daffodil *Narcissus pseudonarcissus*. Engineered biocatalytic methods have the potential to stabilize the supply chain of amaryllidaceae alkaloids. Here, we propose a highly efficient biosensor-AI technology stack for biocatalyst development, which we apply to engineer amaryllidaceae alkaloid production in *Escherichia coli*. Directed evolution is used to develop a highly sensitive (EC_50_= 20 uM) and specific biosensor for the key amaryllidaceae alkaloid branchpoint 4-O’Methylnorbelladine. A machine learning model (MutComputeX) was subsequently developed and used to generate activity-enriched variants of a plant methyltransferase, which were rapidly screened with the biosensor. Functional enzyme variants were identified that yielded a 60% improvement in product titer, 17-fold reduced remnant substrate, and 3-fold lower off-product regioisomer formation.

## Introduction

Alkaloids produced by the *Amaryllidoideae* subfamily of flowering plants have great therapeutic promise, including anticancer, fungicidal, antiviral, and acetylcholinesterase inhibition properties. Among the approximate ∼600 reported AAs, those derived from the lycorine, haemanthamine, and narciclasine scaffolds have been used as lead molecules in anticancer research^1–4^. One of the most notable *Amaryllidoideae* alkaloids (AAs) is galantamine, a selective and reversible acetylcholinesterase inhibitor that is a licensed treatment for mild to moderate symptoms of Alzheimer’s disease and a promising scaffold for drug design^5,6^. Due to galantamine’s challenging synthesis, global supplies largely rely on isolating the low quantities (0.3% dry weight) that accumulate in harvested daffodils, ultimately resulting in an extremely expensive ($50,000/kg) and environmentally-dependent supply chain^7,8^. In an effort to improve galantamine production, new agricultural techniques are currently being tested to boost daffodil-sourced yields^9,10^.

A promising alternative to amaryllidaceae alkaloid extraction from plants is microbial fermentation. Recently, long plant pathways have been reconstituted into microbial hosts for the production of therapeutic benzylisoquinoline alkaloids^11,12^, tropane alkaloids^13^, and monoterpene indole alkaloids^14^. While the complete biosynthetic pathway for any AA with therapeutic value has not yet been elucidated, recent studies have characterized early pathway enzymes responsible for the biosynthesis of 4’-O-Methyl-Norbelladine, the last common intermediate before AA pathway branches diverge^15^. Furthermore, semi-synthetic methods have been proposed using characterized enzymes to generate advanced intermediates^16^. The industrial application of such pathways could be greatly accelerated by augmenting high-throughput screens with genetic biosensors^17–20^, and using artificial intelligence to guide protein design^21–24^, yielding enzymes and pathways with improved stability and activity.

Here, we uniquely synergize the development of custom biosensors with machine learning-guided protein design as a paradigm for rapidly prototyping and improving new pathways. In particular, in order to improve microbial fermentation of the branchpoint AA 4-O’Methylnorbelladine (4NB) a generalist transcription factor, RamR, was evolved into a highly sensitive biosensor for 4NB that precisely discriminates against the non-methylated precursor norbelladine, and the new biosensor was then used to monitor the activity of norbelladine 4-O’Methyltransferase (Nb4OMT) from the daffodil *Narcissus pseudonarcissus* in *Escherichia coli*. A structure-based self-supervised 3D residual neural network (3DResNet) trained to generalize at protein:non-protein interfaces, and the evolved biosensor was used to screen a panel of deep learning-guided Nb4OMT designs. Functional variants of the Nb4OMT enzyme were rapidly identified that yielded a 60% improvement in product titer, 17-fold reduced remnant substrate, and 3-fold lower off-product formation.

## Results

### Identifying a biosensor for the branchpoint amaryllidaceae alkaloid 4-O’Methyl-norbelladine

4-O’Me-norbelladine (4NB) is the branchpoint intermediate for the entire amaryllidaceae alkaloid (AA) family (**Figure 1A**), and therefore was the target compound for biosensor generation. Previously the highly malleable TetR-family *Salmonella typhimurium* repressor RamR had been used as a starting point for identifying biosensors for a variety of benzylisoquinoline alkaloids^17^. Given the chemical similarities between AAs and BIAs, and RamR’s proven ability to rapidly evolve novel ligand specificity, RamR was again used as a starting point for directed evolution.

**Figure 1:**
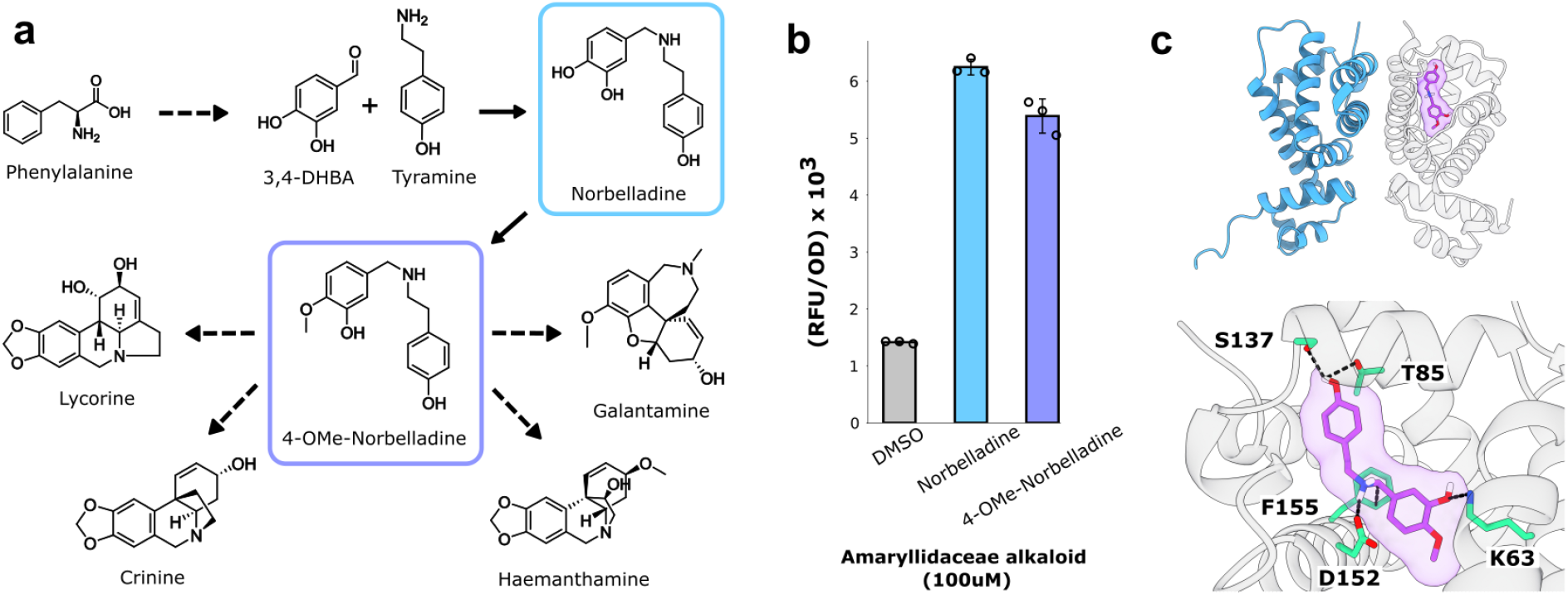
Identifying a biosensor responsive to amaryllidaceae alkaloid intermediates. (**a**). Abbreviated biosynthetic plant pathways for therapeutic amaryllidaceae alkaoids. (**b**) Response of the RamR transcription factor to amaryllidaceae alkaloid pathway intermediates norbelladine and 4-Ome-norbelladine. Error bars represent the S.E.M +/- the mean. Experiments were conducted in biological triplicate.(**c**) Structure of RamR (**PDB: 3VVX**) docked with 4-OMe-norbelladine using GNINA (see **Methods**). Predicted ligand-interacting residues are highlighted green.

The wild-type RamR sensor was constitutively expressed on one plasmid (pReg-RamR) in parallel with another plasmid bearing the regulator’s cognate promoter upstream of the sfGFP gene (Pramr-GFP). Upon induction with various AAs, RamR was found to be slightly responsive to both 4NB and its immediate precursor norbelladine, yielding 3.8 and 4.4-fold increases in fluorescence, respectively (**Figure 1B**). To better understand this promiscuous binding activity, norbelladine was docked within the ligand binding pocket of RamR using GNINA 1.0^25^, and a conformational pose was identified whereby the phenol moiety of norbelladine forms hydrogen bonds with S137 and T85, while the catechol moiety forms a hydrogen bond with K63. This docking pose also suggested that norbelladine’s secondary amine may hydrogen bond with D152 and further interact with the aromatic ring of F155 (**Figure 1C**).

### Evolving a highly specific biosensor for 4-O’Methyl-norbelladine

While the native responsiveness was promising, for practical use in metabolic engineering applications the sensitivity and specificity of RamR for 4NB needed to be greatly improved. The simulated molecular interactions between RamR and 4NB informed a rational approach to library design. Three site-saturated (NNS) RamR libraries that each targeted three residues facing inwards toward the ligand binding cavity were generated (**Supplementary Figure 1**). The 32,000 unique genotypes per library could be readily plumbed using our previously described method, Seamless Enrichment of Ligand Inducible Sensors (SELIS)^17^. Briefly, this method involves a growth-based selection to first filter out biosensor variants that are incapable of repressing transcription from their cognate promoter, followed by a fluorescence-based screen to isolate sensor variants highly responsive to the target analyte.

After the first round of directed evolution, several RamR variants were found to be substantially more responsive to 4NB, even in the absence of a negative selection against norbelladine. In fact, one variant bearing two amino acid substitutions (4NB1.2, K63T and L66M) displayed a 20-fold selectivity for 4NB over norbelladine (**Supplementary Figure 2a, b**). Using 4NB1.2 as a starting point, additional libraries were generated that encompassed the other, previously randomized positions. SELIS was now performed with a growth-based counter-selection against norbelladine (100 uM). The top four biosensor variants were again highly specific for 4NB but now also became significantly more sensitive, with the best variant, 4NB2.1 (C134D and S137G), achieving a limit of detection of approximately 2.5 uM (**Supplementary Figure 2c, d; Figure 2**). Ultimately, the 4NB2.1 sensor was extremely selective for 4NB over norbelladine, displaying an over 80-fold preference for the former, despite the two effectors differing by only a single methyl group.

**Figure 2:**
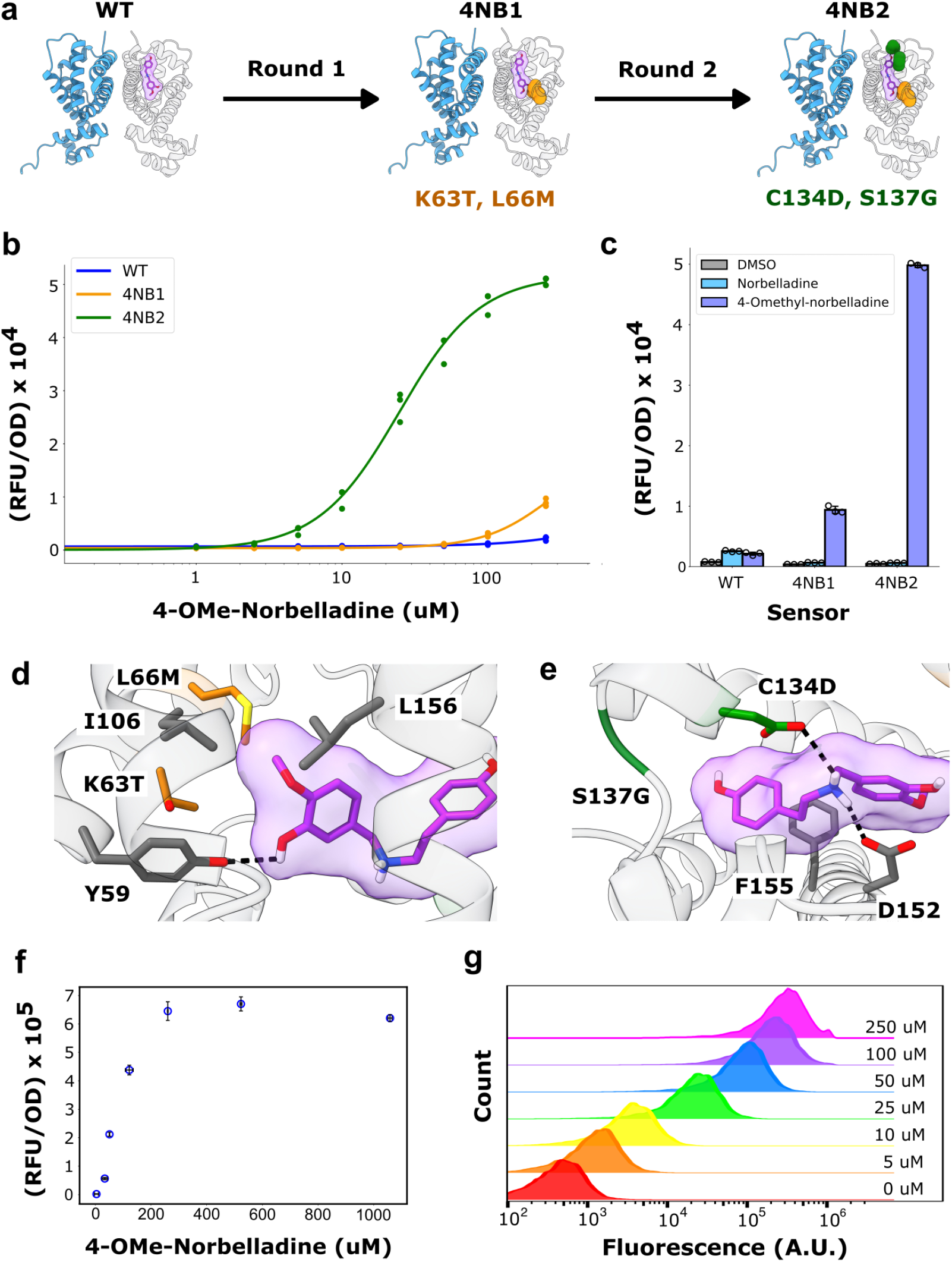
Evolving a highly specific biosensor for 4’-O-Methyl-Norbelladine. (**a**) Schematic illustrating the mutations that resulted after round one (4NB1) and round two (4NB2) of RamR evolution towards 4-OMe-norbelladine. (**b**) Dose response measurements of WT RamR, 4NB1, and 4NB2 mutants with 4-OMe-norbelladine. (**c**) Relative response of WT RamR, 4NB1, and 4NB2 mutants to norbelladine and 4-Ome-norbelladine. (**d, e**) Structure of Alpha Folded 4NB2 docked with 4-OMe-norbelladine. Predicted ligand interactions with WT residues, mutations that arose in 4NB1, and mutations that arose in 4NB2 are colored gray, orange, and green, respectively. (**f**) Correlation between fluorescent response measured with the 4NB2 sensor and 4-OMe-norbelladine measured with HPLC. (**g**) The distribution of fluorescent cell populations in response to 4-OMe-norbelladine concentration. All data was performed in biological triplicate. Error bars represent the S.E.M +/- the mean.

To again explore the structural basis for precise methyl group discrimination a structural model of 4NB2.1 was generated using AlphaFold 2.0^26^, and 4NB was docked into this model using GNINA 1.0^25^. The docked pose suggests that the K63T substitution repositions the hydroxyl group at position 3 of 4NB to hydrogen bond with the wild-type Y59 residue, while the L66M substitution strengthens a hydrophobic pocket around the 4-O’Methyl group of 4NB (along with the native I106 and L156 residues; **Figure 2d**). This analysis is in agreement with the fluorescence assay data, since only RamR variants bearing the K63T and L66M mutations are highly selective for 4NB over norbelladine (**Supplementary Figure 2**). The model also positions the new aspartate at position 134 (C134D) to hydrogen bond with the amine of 4NB; several other RamR variants also placed a hydrogen bond donor (glutamate, glutamine, asparagine) at the 134 position (**Supplementary Figure 2**). Overall, as a consequence of these substitutions the 4NB ligand may shift in position to allow for more favorable pi-pi stacking with F155 (**Figure 2e**).

To evaluate the utility of the 4NB2.1 sensor for high-throughput screening of AAs, we compared its performance to an HPLC method adapted from the literature^27^. The concentration range of 4NB can be discerned between 2.5 uM and 250 uM, while the equivalent range for the HPLC method is between 25 uM and 1000 uM (**Figure 2f**). The dynamic range of sensing could potentially be further increased via less sensitive biosensor intermediates characterized during evolution (see **Supplementary Figure 2**).

Most importantly, the 4NB2.1 sensor is approximately 10-fold more sensitive than the HPLC method, making it well-suited for screening transplanted biosynthetic enzymes from plants, which often initially show low flux^28^. Flow cytometry analysis indicated that the sensor’s response at the population level was highly uniform (**Figure 2g**), ensuring low noise measurements.

### Monitoring norbelladine O-methyltransferase activity in Escherichia coli

Although several AAs have been recognized for their therapeutic value, to our knowledge there have so far been no attempts to reconstitute AA pathways in microbial hosts. Since norbelladine 4-O-methyltransferase (Nb4OMT) from the wild daffodil *Narcissus sp. aff. pseudonarcissus*, is directly responsible for 4NB production from norbelladine, we chose this as a starting point for development of a fuller pathway. A 4NB reporter plasmid (pSens4NB2; **Supplementary Figure 3**) was co-transformed with a plasmid constitutively expressing Nb4OMT. When this strain was grown in media supplemented with the substrate norbelladine, Nb4OMT activity could be observed, monitored, and quantified via fluorescence (**Figure 3a**). The level of cell fluorescence correlated positively with enzyme expression strength (**Supplementary Figure 4**), with the concentration of norbelladine supplemented into the culture media, and with 4NB titer measured via HPLC (**Figure 3b**). As was the case with measuring 4NB supplemented media, the fluorescence of cellular populations was uniformly distributed, again indicating that there was little noise during production or sensing. The independent measurements of noise via the 4NB biosensor will likely prove important as high yield strains are further developed and translated.

**Figure 3.**
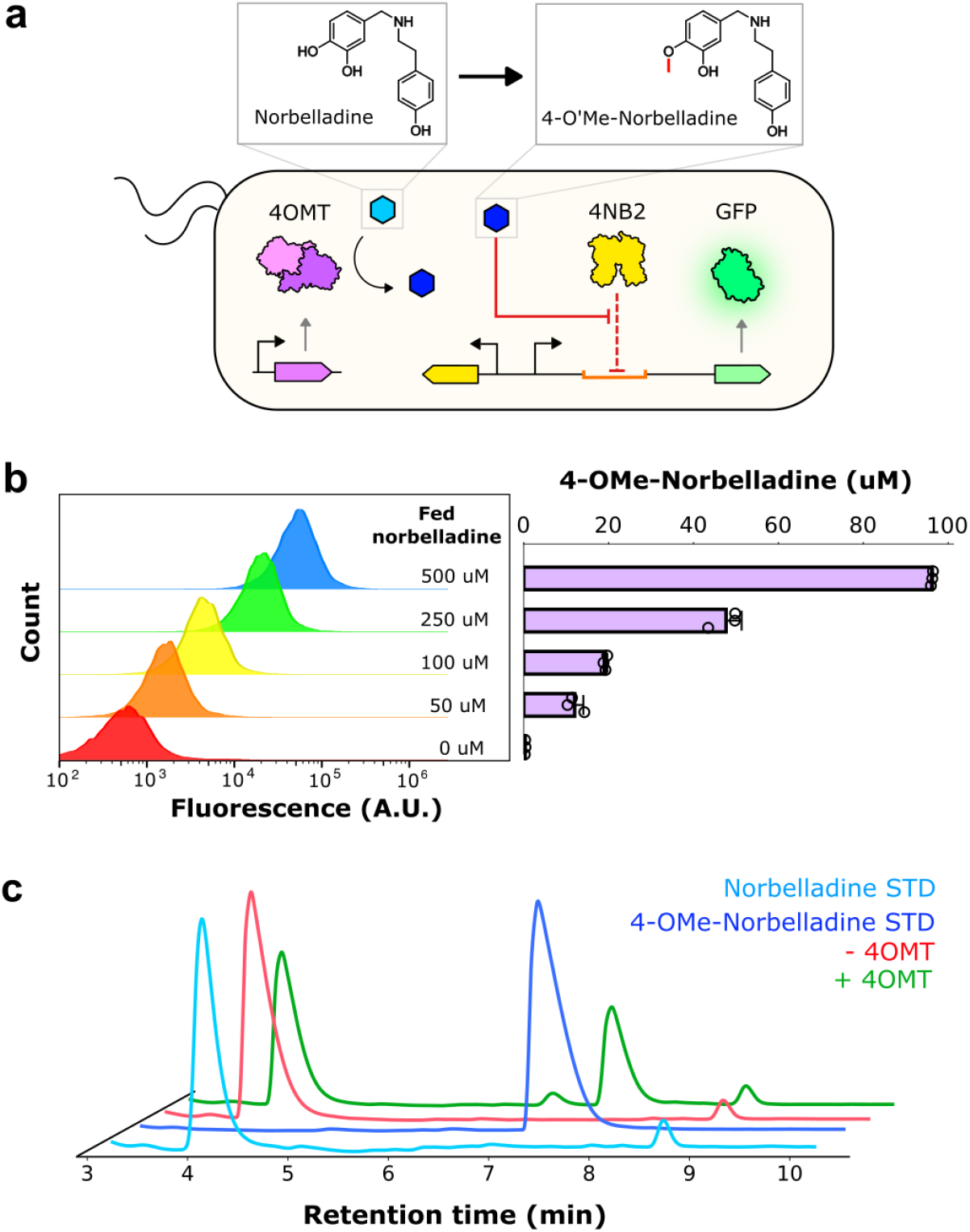
Monitoring 4OMT activity with the 4NB2 biosensor. (**a**) Schematic representation of the biosensor-monitored enzymatic reaction within E. coli cells. Light blue and dark blue hexagons denote norbelladine and 4-OMe-norbelladine, respectively. (**b**) Correlation between cell population fluorescence and biosynthesized 4-OMe-norbelladine, measured by HPLC. Error bars represent S.E.M. +/- the mean. (**c**) Chromatographic traces of supernatant collected from *E. coli* cells expressing an active Nb4OMT enzyme (green) or an empty plasmid (red). Traces for norbelladine and 4-OMe-norbelladine standards are shown in light blue and dark blue, respectively.

While these results demonstrated the utility of the evolved biosensor for monitoring Nb4OMT activity, they also revealed the catalytic inefficiency of the enzyme. HPLC analysis indicated that a significant amount of supplemented norbelladine remained after culturing for 24 hours (**Figure 3c**). Indeed, leftover norbelladine was identified when as low as 50 uM of norbelladine was supplemented into the culture media (**Supplementary Figure 5**). Furthermore, LC/MS analysis identified 3-OMe-norbelladine as a minor component, indicating that the wild-type Nb4OMT enzyme was not highly regiospecific (**Supplementary Figure 6**). These observations all suggested that Nb4OMT activity and specificity could be improved by enzyme engineering.

### Combining custom biosensors and machine learning to improve pathway activity

To improve Nb4OMT activity in a microbial host we initially carried out directed evolution starting from randomly mutagenized libraries, via error-prone PCR. The library of enzyme variants was transformed into cells containing the pSens4NB2.1 plasmid, plated on solid media containing norbelladine, and highly-fluorescent colonies were isolated and then individually phenotyped in a secondary liquid-based fluorescence screen. Interestingly, while this approach had previously proven effective for identifying improved enzyme variants in other pathways^17^, it failed to enhance Nb4OMT activity.

We therefore pursued a complementary approach to enzyme engineering, using machine learning to better identify variants and potential library designs. A structure-based convolutional neural network (CNN; MutCompute) had previously proven adept at predicting mutations that improved protein functionalities, including fluorescence (BFP)^29^, expression (PMI)^29^, stability (polymerase)^30^, and catalytic activity (PETase)^21^. Unfortunately, the structure of the Nb4OMT enzyme had not been solved, preventing the generation of structure-based CNN predictions for substitutions. Instead, a *de novo* structural model for Nb4OMT was generated using Alphafold2^26^, and both the S-adenosyl-homocysteine (SAH) cofactor and norbelladine were docked using GNINA1.0^25^. The SAH cofactor was chosen instead of SAM because the nearest structure, of Alfalfa caffeoyl coenzyme A 3O-methyltransferase (PDB: 1SUI; sequence similarity: 60.79%), contained this cofactor, and its SAH pose was transplanted to the AlphaFolded Nb4OMT scaffold. GNINA scored the minimized SAH pose with a 0.835 probability of being within 2Å RMSD from the real pose, and predicted an affinity of -7.9 kcal/mol (**Supplementary Table 1**). The GNINA pose was guided by the supposition that either D155 or K158 must be the general-base that deprotonates the 4-hydroxyl group during the SN2 reaction, and that a potential cation-pi interaction with K158 would orient the plane of the catechol ring in the active site. GNINA scored the minimized norbelladine pose with a 0.824 probability of being within 2Å RMSD from the real pose and a predicted affinity of -7.3 kcal/mol (**Supplementary Table 1**).

The original data engineering pipelines established for MutCompute restricted its training to microenvironments with atoms belonging to the 20 amino acids, and therefore MutCompute was unable to provide contextualized predictions in microenvironments that possessed atoms from cofactors or ligands^29,31^. To address this, we 1) rebuilt the data engineering pipelines to enable training on heterogenous microenvironments (see **Methods**), 2) curated new training and testing datasets that prioritized sampling these heterogeneous microenvironments (see **Methods**), and 3) developed a novel residual convolutional architecture to improve feature extraction capabilities and in turn the predictive power of the model (**Supplementary Figure 7**). The new self-supervised 3D residual neural network (3DResNet) achieved improved wildtype prediction accuracy ∼80% on a ∼250K residue test set compared to 69% on a 6K test set from the previous 3DCNN model^29,31^. Furthermore, the 3DResNets were shown to generalize to protein-ligand interaction interfaces without any drop in wildtype accuracy (81% wildtype accuracy on a protein-ligand interface test set compared to 62.1% from the previous 3DCNN model). After training numerous models, we selected models for ensembling and ml-engineering of norbelladine methlytransferase based on their zero-shot capability to correlate with ΔTM point mutations from FireProtDB^32^ (zero-shot correlation described in methods). The ensembled 3DresNet model had an overall wildtype accuracy of 67.3% and protein-ligand interface wildtype accuracy of 66.%. The ternary computational Nb4OMT docked complex was passed to the improved 3DResNet model, and predictions were generated for each amino acid throughout the Nb4OMT protein (**Figure 4a**). Based on these predictions, we manually curated predicted substitutions, prioritizing those that were near the active site and that were likely to form known stabilizing motifs such as salt bridges.

**Figure 4.**
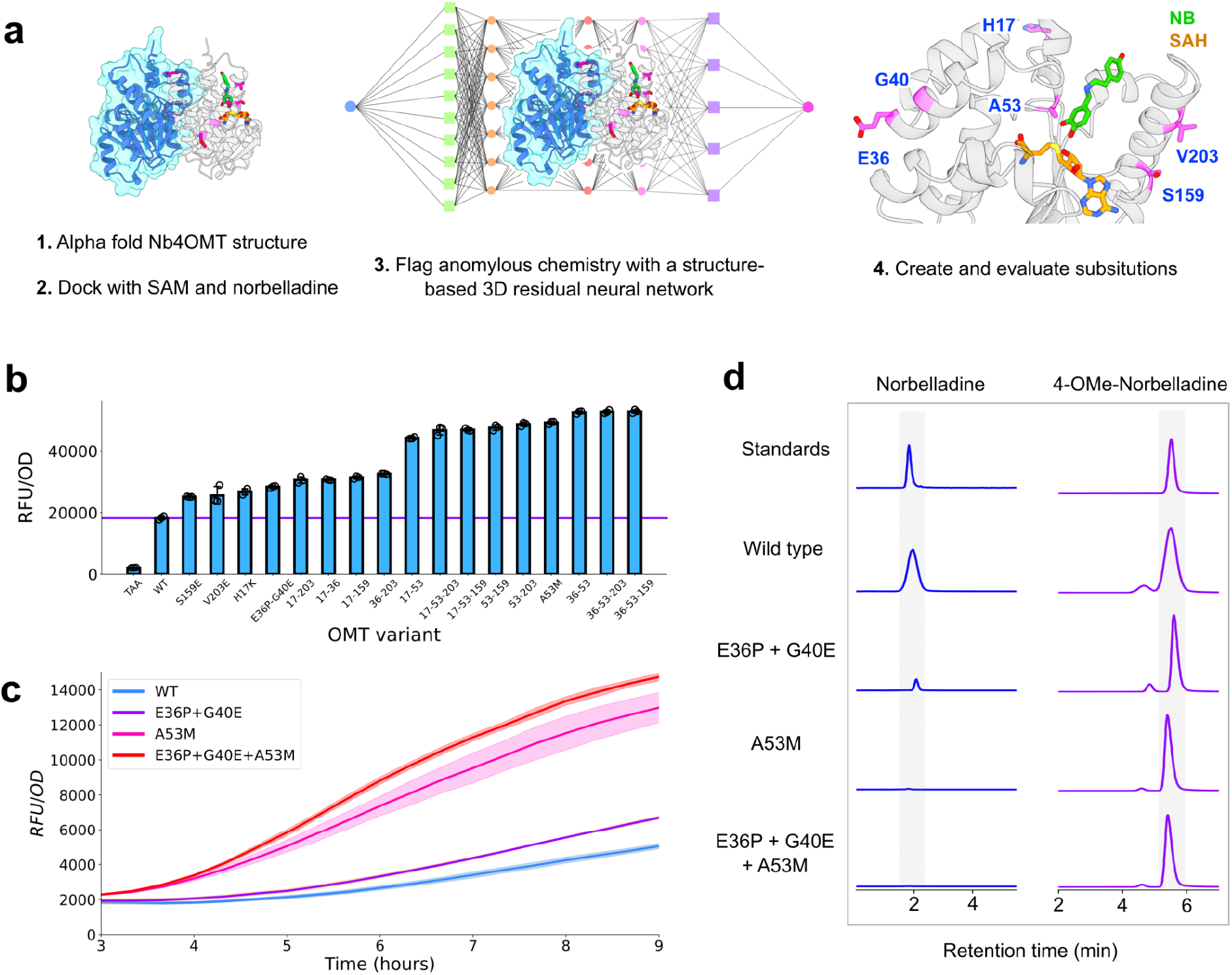
Synergizing AI and biosensors to improve 4NB titers. (**a**) MutComputeX workflow for generating AI-guided enzyme designs. (**b**) Fluorescent signal produced from *E. coli* cells containing the 4-OMe-norbelladine reporter plasmid (pSens-4NB2) and expressing either an empty plasmid (TAA), the wild-type Nb4OMT enzyme (WT), or AI-designed Nb4OMT mutants, when cultured with 100 uM of norbelladine at 37°C. The purple horizontal line denotes the fluorescent signal produced from culturing the wild-type Nb4OMT enzyme. Error bars represent the S.E.M. +/- the mean (**c**) Time-dependent fluorescent signals produced by *E. coli* cells containing the 4-OMe-norbelladine reporter plasmid (pSens-4NB2) and expressing Nb4OMT (WT) or AI-designed mutants. (**d**) Ion-extracted chromatograms of chemical standards (blue) or the supernatant of cells expressing Nb4OMT or AI-designed mutants (purple) cultured with norbelladine.

Ultimately, 22 mutational designs were experimentally validated in *E. coli*. Leveraging the biosensor-enabled high-throughput screen, we were able to quickly assess each of the 22 mutants across three temperatures (25°C, 30°C, 37°C) and two substrate concentrations (100 uM, 1 mM). In all tested conditions, the A53M mutation consistently produced a fluorescent signal significantly above the wild-type enzyme, while the H17K, H17R, S159E, V203E, and E36P-G40E substitutions produced signals above wild-type in at least one tested condition (**Supplementary Figure 8**). Increasing the reaction temperature to 37°C improved product formation (despite the fact that the *Narcissus pseudonarcissus* plant grows in 10-23°C climates^33^). Double and triple mutants incorporating the H17K, A53M, S159E, V203E, and E36P-G40E substitutions were generated and screened; as with the initial screens, variants bearing the A53M mutation produced the greatest signals (**Figure 4b**). In time course reactions, after media supplementation with norbelladine the rate of fluorescence increase for the E36P-G40E variant was similar to that of the wild-type enzyme, but the rates produced by the two A53M-bearing variants were significantly higher (**Figure 4c**). LC/MS analyses were carried out on supernatants from the E36P-G40E, A53M, and E36P-G40E-A53M variants, and in agreement with our fluorescence-based assay the level of 4NB product increased by 60% while the level of remnant norbelladine decreased 17-fold (**Figure 4d**, **Supplementary Figure 9**). The A53M mutation reduced levels of the 3-O’Methyl-norbelladine off-product by about 3-fold (**Supplementary Figure 10)**.

## Discussion

Herein we report the use of directed evolution and machine learning-guided design for the development of custom microbial biosensors that could be used to monitor substantive improvements in amaryllidaceae alkaloid pathway activity. The RamR transcription factor was evolved to respond to low micromolar levels of the pathway branchpoint 4NB, and after only four substitutions exquisite specificity emerges for the methylated oxygen moiety in 4NB, with a barely detectable response to the non-methylated precursor norbelladine. Overall, these results highlight the powerful capability of using evolved biosensors for precisely reporting on pathway intermediates while avoiding cross-reactivity with closely related precursor molecules. The RamR protein is now well positioned as an ideal starting point for the generation of biosensors for not only benzylisoquinoline alkaloids, but also for AAs such as galantamine, haemanthamine, lycorine, and their intermediates.

The high specificity was also essential for measuring the real-time activity of the plant-derived Nb4OMT enzyme in *E. coli*, which in turn allowed us to leverage our state-of-the-art 3DResNet, MutComputeX for enzyme engineering. MutComputeX was trained on ∼2.3M microenvironments sampled from over 23,000 protein structures, and predicted functional variants of the Nb4OMT enzyme with 60% improvement in product titer, 17-fold reduced remnant substrate, and 3-fold lower off-product formation. Starting for the first time from an AlphaFold structure model docked with its substrate and cofactor, MutComputeX designs yielded variants with not only improved product:substrate ratios, but also improved regiospecificities, as determined by LC/MS analysis.

That said, many challenges remain to realizing a commercially viable microbial strain for the fermentation of AAs. First, the central precursors 3,4-dihydroxybenzoate and tyramine must be overproduced in a base strain, likely *Saccharomyces cerevisiae* due to its proven ability to functionally express multiple plant-derived cytochrome proteins ^34^. Biosynthetic pathways for the production of 3,4-dihydroxybenzoate and tyramine have already been engineered into *S. cerevisiae* ^35,36^, with high-throughput screening methods yielding improved tyrosine precursor yields ^37^. Second, the early pathway enzymes for the production of norbelladine, norcraugsodine reductase, and norbelladine synthase must be functionally expressed in a microbial strain. So far, the activities of these enzymes have only been demonstrated within an *in vitro* context ^38–40^. Third, the downstream enzymes necessary for the production of advanced AAs must be identified and functionally expressed. With regards to galantamine, the CYP96T1 cytochrome protein has been shown to catalyze the *para-ortho* coupling of 4NB and thus produce the galantamine precursor N-demethylnarwedine, but only as a minor product making up only 1% of the final product^41^. Finally, the remaining N-methyltransferase and ketone reductase enzymes necessary for complete galantamine biosynthesis remain unknown, although transcriptomic datasets from several daffodil species exist and comparative transcriptomic analyses have proven successful in the past for identifying missing pathway enzymes ^15^.

While the development of strains for therapeutic AA biosynthesis is a significant endeavor, we believe that the approach presented in this work, synergizing custom biosensor-enabled screens with self-supervised machine learning-guided protein design, will fundamentally accelerate the pace of strain and enzyme engineering as a whole. Custom biosensor-enabled screens enable rapid collection of phenotype data under a wide variety of experimental conditions, including determining the kinetics of product formation among strain and enzyme variants, values that are nearly impossible to measure using traditional analytical instruments. The importance of machine learning is further highlighted by failed attempts to engineer Nb4OMT using random mutagenesis alone. Into the future, microbial semi-synthesis of galantamine and other AAs could provide faster production cycles, a more reliable supply chain, and reduced land and water use compared to traditional plant harvesting methods, and the biosensor-AI hybrid technology stack we have advanced herein should greatly accelerate the engineering of upstream enzymes in the pathway, such as norbelladine synthase and norcraugsodine reductase ^38,40^.

## Methods

### Strains, plasmids and media

*E. coli* DH10B (New England Biolabs) was used for all routine cloning and directed evolution. All biosensor systems were characterized in *E. coli* DH10B. LB Miller (LB) medium (BD) was used for routine cloning, fluorescence assays, directed evolution and orthogonality assays unless specifically noted. LB with 1.5% agar (BD) plates were used for routine cloning and directed evolution. The plasmids described in this work were constructed using Gibson assembly and standard molecular biology techniques. Synthetic genes, obtained as gBlocks, and primers were purchased from IDT. Plasmid designs and sequences are listed in **Supplementary Table 4.**

### Chemicals

4-O’Methylnorbelladine was purchased from Toronto Research Chemicals (Toronto Research Chemicals. CAT#: H948930). Tyramine (T90344), 3,4-dihydroxybenzaldehyde (37520), dichloromethane (439223), and NaBH_4_ were purchased from Sigma Aldrich. NMR solvents (d^6^-DMSO, CD_3_OD) were purchased from Cambridge isotope laboratories.

### Chemical synthesis and NMR analysis of norbelladine

The aldehyde (3,4-dihydroxybenzaldehyde) (1 mM, 138 mg) and tyramine (1 mM, 137 mg) were dissolved in dichloromethane (5 mL) and converted to the imine in situ compound by stirring for 4 hr at room temperature. The imine compound was reduced with NaBH_4_ (2 mM, 75.6 mg), washed with water and dried to produce crude product. The crude material was then purified by combinatorial flash chromatography to yield norbelladine (10-90% MeCN in H_2_O, 20 min; 130 mg recovered, beige orange solid, 50 % yield), which was confirmed via NMR (**Supplementary Figure 10**). NMR spectra were taken on the 500 MHZ Bruker prodigy at University of Texas at Austin.

### Chemical transformation

For routine transformations, strains were made competent for chemical transformation. Five milliliters of an overnight culture of DH10B cells was subcultured into 500 mL LB medium and grown at 37 °C and 250 r.p.m. until an optical density of 0.7 was reached (∼ 3 h). Cultures were centrifuged (3,500g, 4 °C, 10 min), and pellets were washed with 70 mL chemical competence buffer (10% glycerol, 100 mM CaCl2) and centrifuged again (3,500g, 4 °C, 10 min). The resulting pellets were resuspended in 20 mL chemical competence buffer. After 30 min on ice, cells were divided into 250-μL aliquots and flash frozen in liquid nitrogen. Competent cells were stored at −80 °C until use.

### Biosensor response assay

The pReg-RamR and Pramr-GFP plasmids were co-transformed into DH10B cells, which were then plated on LB agar plates containing appropriate antibiotics. Three separate colonies were picked for each transformation and were grown overnight. The following day, 20 μL of each culture was then used to inoculate six separate wells in a 2-mL 96-deep-well plate (Corning, P-DW-20-C-S) sealed with an AeraSeal film (Excel Scientific) containing 900 μL LB medium, one for each test ligand and a solvent control. After 2 h of growth at 37 °C, cultures were induced with 100 µL LB medium containing either 10 μL DMSO or 100 μL LB medium containing the target AA dissolved in 10 μL DMSO. Cultures were grown for an additional 4 h at 37 °C and 250 r.p.m. and subsequently centrifuged (3,500g, 4 °C, 10 min). Supernatant was removed, and cell pellets were resuspended in 1 mL PBS (137 mM NaCl, 2.7 mM KCl, 10 mM Na2HPO4, 1.8 mM KH2PO4, pH 7.4). For bulk fluorescence measurements, one hundred microliters of the cell resuspension for each condition was transferred to a 96-well microtiter plate (Corning, 3904), from which the fluorescence (excitation, 485 nm; emission, 509 nm) and absorbance (600 nm) were measured using the Tecan Infinite M1000 plate reader. For single cell fluorescence measurements, 20 uL of cells were diluted in 1 mL of PBS within a clear 96-well microtiter plate and analyzed on the Attune NxT flow cytometer (Thermo Fisher).

### RamR library design and construction

Three semi-rational libraries were designed, each targeting three inward-facing residues within the RamR ligand-binding pocket (**Supplementary Figure 1**). Libraries were generated using overlap PCR with redundant NNS codons using AccuPrime Pfx (Thermo Fisher, 12344024) and cloned into pReg-RamR. *E. coli* DH10B bearing pSELIS-RamR was transformed with the resulting library. Transformation efficiency always exceeded 10^6^ for each round of selection, indicating several fold coverage of the library. Transformed cells were grown in LB medium overnight at 37 °C with carbenicillin and chloramphenicol.

### Directed evolution of RamR biosensors

Cell culture (20 μl) bearing the sensor library was seeded into 5 ml fresh LB containing appropriate antibiotics, 100 μg ml−1 zeocin (Thermo Fisher, R25001) and 100 μM of norbelladine (for round two) and grown at 37 °C for 7 h. Following incubation, 0.5 μl of culture was diluted into 1 ml LB medium, from which 100 μl was further diluted into 900 μl LB medium. Three hundred microliters of this mixture was then plated across three LB agar plates (100 uL per plate) containing carbenicillin, chloramphenicol and 4NB dissolved in DMSO. Plates were incubated overnight at 37 °C. The following day, the brightest colonies were picked and grown overnight in 1 ml LB medium containing appropriate antibiotics in a 96-deep-well plate sealed with an AeraSeal film at 37 °C. A glycerol stock of cells containing **pSELIS-RamR** and pReg-RamR encoding the template RamR variant was also inoculated into 5 ml LB for overnight growth.

The following day, 20 μl of each culture was used to inoculate two separate wells in a new 96-deep-well plate containing 900 μl LB medium. Additionally, eight separate wells containing 1 ml LB medium were inoculated with 20 μl of the overnight culture expressing the parental RamR variant. After 2 h of growth at 37 °C, the top half of the 96-well plate was induced with 100 μl LB medium containing 10 µl DMSO, whereas the bottom half of the plate was induced with 100 μl LB medium containing 4NB dissolved in 10 μl DMSO. The concentration of 4NB used for induction is typically the same concentration used in the LB agar plate for screening during that particular round of evolution. Cultures were grown for an additional 4 h at 37 °C and 250 r.p.m. and subsequently centrifuged (3,500g, 4 °C, 10 min). Supernatant was removed, and cell pellets were resuspended in 1 ml PBS. One hundred microliters of the cell resuspension for each condition was transferred to a 96-well microtiter plate, from which the fluorescence (excitation, 485 nm; emission, 509 nm) and absorbance (600 nm) were measured using the Tecan Infinite M1000 plate reader. Clones with the highest signal-to-noise ratio (generally the top 5–10% of the screened clones) were then sequenced and subcloned into a fresh pReg-RamR vector.

For sensor variant validation, the subcloned pReg-RamR vectors expressing the sensor variants were transformed into DH10B cells expressing Pramr-GFP. These cultures were then assayed, as described in **Biosensor response assay**, using eight different concentrations of the 4NB. The sensor variant that displayed a combination of low background, a reduced EC_50_ for 4NB and a high signal-to-noise ratio was then used as the template for the next round of evolution.

### Dose–response measurements

Glycerol stocks (20% glycerol) of strains containing the plasmids of interest were inoculated into 1 ml LB medium and grown overnight at 37 °C. Twenty microliters of overnight culture was seeded into 900 μl LB medium containing ampicillin and chloramphenicol in a 2-ml 96-deep-well plate sealed with an AeraSeal film. Following growth at 37 °C and 250 r.p.m. for 2 h, cultures were induced with 100 μl of an LB medium solution containing appropriate antibiotics and the inducer molecule dissolved in 10 μl DMSO. Cultures were grown for an additional 4 h at 37 °C and 250 r.p.m. and subsequently centrifuged (3,500g, 4 °C, 10 min). Supernatant was removed, and cell pellets were resuspended in 1 ml PBS. The cell resuspension (100 μl) for each condition was transferred to a 96-well microtiter plate, from which the fluorescence (excitation, 485 nm; emission, 509 nm) and absorbance (600 nm) were measured using the Tecan Infinite M1000 plate reader.

### Biosensor-linked O-methyltransferase activity assay

Nb4OMT was expressed with the P150-RBS(riboJ) promoter–RBS on the pReg-RamR plasmid backbone (no regulator present). Cells were co-transformed with both the 4OMT plasmid and the 4NB reporter plasmid and plated on an LB agar plate containing appropriate antibiotics. Three individual colonies from each transformation were picked into LB and grown overnight. Resulting cultures were diluted 50-fold into 1 ml LB medium containing the indicated concentration of norbelladine in a 96-deep-well plate and were grown at the indicated temperature for 24 h. Subsequently, the fluorescence of cultures was measured in the same manner as previously described in Dose–response measurement above.

### Ternary Complex Generation with AlphaFold2 and GNINA1.0

Nb4OMT wild type sequence (uniprot id: A0A077EWA5) was run through the AlphaFold2-multimer as a homodimer using the publicly available collab notebook. This resulted in a computational structure with a pLDDT of 0.955 and a pTM of 0.94. The initial coordinates for the SAH cofactor were transplanted onto the AlphaFold structure from the 1SUI pdb structure and then optimized with GNINA1.0’s –local_only and –minimize flags. Norbelladine’s initial 3D coordinates were obtained from the PubChem database (id: 416247) and docked into the active site of the A protomer. To dock norbelladine, we generated a bounding box for the GNINA docking procedure by finding the largest 3D box from the atomic coordinates of the following residues: L10, W50, S52, A53, D155, D157, K158, W185, Y186, A204. GNINA was run several times with different seeds and all docked poses were manually screened for known mechanistic insight. The docked pose that best satisfied the mechanistic insight and received a high GNINA docking score was then minimized with the –local_only and –minimize flags. The docking results from GNINA for SAH and NB are shown in **Supplementary Table 1**.

### Building MutComputeX

#### Structure File pre-processing

To generate voxelized matrices of microenvironments that span between protein:non-protein atoms, experimental CIF files were pre-processed with 1) ChimeraX to add hydrogen atoms to the proteins, nucleic acids, and organic ligands; 2) ChargeFW2 to add polarized charges that bridge protein: non-protein interfaces; and 3) FreeSASA to add solvent accessible surface area values that take into account protein:non-protein interactions. CIF read and write functionality for ChargeFW2 and FreeSASA were implemented and merged to both open-sourced libraries.

#### Voxelized Matrix Generation

To generate a voxelized molecular representation of a microenvironment, a 20Å cube of atoms was filtered from the structure centered on the Calpha and oriented with respect to the backbone where the side chain was along the +z axis. All atoms in the center residue are then removed prior to insertion into a voxelized grid with 1Å resolution. Each atom is placed into a corresponding element channel except halogen atoms (which are placed into a multi-atom channel that consist of F, Cl, Br, I) and each atom’s partial charge and SASA value are placed into the partial charge and SASA channels, respectively. For all channels, atom values are gaussian blurred according to their Van-der-Waals radii.

#### Dataset Generation

A dataset of 50% sequence similar protein chains with at least a 3.0Å resolution was downloaded in November 2021 from the RCSB. This provided us with X protein sequences from Y PDB entries. To generate microenvironment datasets, each protein where residues within 5Å of a non-protein entity were prioritized and then randomly backfilled until 200 residues or half of the protein sequence was sampled. A total of 2,569,256 microenvironments were sampled from 22,759 protein sequences and split 90:10 to generate our training and test set splits for interfaces and non-interface residues are shown in **Supplementary Table 2**.

#### Model Training

The 3D residual neural network was built in Tensorflow 2.x. The architecture is provided in **Supplementary Figure 7**. Each model run was parallelized over 4 AMD Radeon Instinct MI50s with a batch size of 200. Models were trained for up to 8 epochs where each epoch was saved as a checkpoint with a variety of hyperparameters. We used a scheduled learning rate that began at 0.001 and had an exponential decay constant of either 0.3 or 0.5 and an adaptive learning rate that would lower the learning rate by 0.25 if the training accuracy did not improve by 0.1% after either 30K, 50K, and 60K training instances. Weights were updated with the Adam optimizer and all convolutional layers had weight decay regularization of 0.001.

#### Model Benchmarking

To ensure our datasets were enabling the 3D ResNet models to generalize across protein: non-protein interfaces, we monitored the overall wildtype accuracy and wildtype accuracy for residues at DNA, RNA, and ligand interfaces on our test set. To select models to ensemble and generate engineering predictions we generated zero shot-predictions for all mutational data in FireProtDB and chose the models that had the highest correlation with the single point mutation ΔTM experimental data. The zero-shot predictions were generated by taking the prediction assigned to the wildtype and mutant amino acid from FireProtDB and taking the log odds where a positive log odd means a stabilizing prediction and a negative log odd means a destabilizing prediction. The ensembled model had a Pearson and Spearman correlation coefficients of 0.367 and 0.425 with the 2719 single point mutations with ΔTM experimental data in FireProtDB and a Pearson and Spearman correlation coefficients of -0.407 and -0.457 with the 4889 single point mutations with ΔΔG experimental data in FireProtDB. Correlation coefficients for the independent models can be found in **Supplementary Table 3.**

### Mutational Designs

Mutations were designed with two goals: stabilizing the protein away from the active site and investigating point mutations where predictions differed between the docked and apo protein structures. With these objectives, we sorted residues based on the log odds between the predicted and wild type amino acids. For the stability objective, predictions that recapitulate known chemical phenomena such as salt bridges, hydrogen bonding, proline capping were prioritized.

### High-performance liquid chromatography analysis

Assay samples were filtered using a 0.2-um PTFE syringe filter prior to running the HPLC. The measurement of Norbelladine and 4-O’Methyl-norbelladine was performed using a Vanquish HPLC system (Thermo Fisher Scientific) equipped with a BDS Hypersil TM C18 (3.0×150mm 2, 3um) (Thermo Fisher Scientific) with detection wavelength 277 nm. The mobile phase consisted of 0.1% formic acid in water or 0.1% formic acid in acetonitrile over the course of 28 minutes under the following conditions: 10% organic (vol/vol) for 2 minutes, 10 to 30% organic (vol/vol) for 13 minutes, 30 to 90% organic (vol/vol) for 0.1 minutes, 90% organic (vol/vol) for 4.9 minutes, 90 to 10% organic (vol/vol) for 1 minute, and 10% organic (vol/vol) for 7 minutes. The flow rate was fixed at 0.8 ml min -1. A standard curve for norbelladine was prepared using synthesized norbelladine (see **Chemical synthesis and NMR analysis of norbelladine**). A standard curve for 4-O’Methyl-norbelladine was prepared using commercially available 4-O’Methyl-norbelladine.

### Liquid chromatography–mass spectrometry

Cells containing the plasmid expressing each Nb4OMT variant with the P150-RBS(RiboJ) promoter were transformed and plated onto an LB agar plate containing appropriate antibiotics. The following day, three colonies from each plate were cultured overnight in LB and subsequently diluted 50-fold into 1 ml LB containing 1 mM norbelladine. These cultures were grown for 24 h at 37 °C and centrifuged at 16,000g for 1 min, and the resulting supernatant was filtered using a 0.2-μm filter.

Samples were analyzed using an Agilent 6530 Q-TOF LC–MS with a dual Agilent Jet Stream electrospray ionization source in positive mode. Chromatographic separations were obtained under gradient conditions by injecting 10 μl onto an Agilent RRHD Eclipse Plus C18 column (50 × 2.1 mm, 1.8- μm particle size) with an Agilent ZORBAX Eclipse Plus C18 narrow-bore guard column (12.5 × 2.1 mm, 5-μm particle size) on an Agilent 1260 Infinity II liquid chromatography system. The mobile phase consisted of eluent A (water with 0.1% formic acid) and eluent B (acetonitrile). The gradient was as follows: Hold 95% A / 5% B from 0 to 2 min (0.7 ml min−1), 80% A / 20% B from 2 to 15 min (0.7 ml min−1), 70% A / 5% B from 15 to 18 min (0.7 ml min−1). The sample tray and column compartment were set to 7 °C and 30 °C, respectively. The fragmentor was set to 100V. Q-TOF data were processed using the Agilent MassHunter Qualitative Analysis software. Both products and the residual substrate of the wildtype reactions were identified with MS/MS with a collision cell energy of 5 V. To create the chromatograms (shown in **Fig. 4C** and **Supplementary Figure 6**), signal counts from the EIC within a window ±0.05 min relative to the retention time of the substrate and products were extracted for each scan (m/z ratios 260.1281 and 274.1438).

### Statistical analysis and reproducibility

All data in the text are displayed as mean ± s.e.m. unless specifically indicated. Bar graphs, fluorescence and growth curves, dose–response functions were all plotted in Python 3.6.9 using Matplotlib. Dose–response curves and EC_50_ values were estimated by fitting to the Hill equation y = d + (a − d)xb(cb + xb)−1 (where y = output signal, b = Hill coefficient, x = ligand concentration, d = background signal, a = maximum signal and c = EC_50_), with the scipy.optimize.curve_fit library in Python.

## Data availability

Protein sequence information was retrieved from the NCBI database: RamR, 3VVX_A; Nb4OMT, A0A077EWA5.1. Plasmid sequences relevant to this study are positioned for deposition in Addgene.

## Code availability

Code used to generate bar plots and dose–response functions presented in this text is accessible at https://github.com/simonsnitz/plotting. The MutComputeX model as well as the input data of the norbelladine-4O-methyltransferase are available at https://github.com/danny305/MutComputeX.

## References

1. Berkov, S., Osorio, E., Viladomat, F. & Bastida, J. Chapter Two - Chemodiversity, chemotaxonomy and chemoecology of Amaryllidaceae alkaloids. in The Alkaloids: Chemistry and Biology (ed. Knölker, H.-J.) vol. 83 113–185 (Academic Press, 2020).

2. Evidente, A. et al. Biological Evaluation of Structurally Diverse Amaryllidaceae Alkaloids and their Synthetic Derivatives: Discovery of Novel Leads for Anticancer Drug Design. Planta Med. 75, 501–507 (2009).

3. Cahlíková, L. et al. The Amaryllidaceae alkaloids haemanthamine, haemanthidine and their semisynthetic derivatives as potential drugs. Phytochem. Rev. 20, 303–323 (2021).

4. Roy, M. et al. Lycorine: A prospective natural lead for anticancer drug discovery. Biomed. Pharmacother. 107, 615–624 (2018).

5. Bhattacharya, S., Maelicke, A. & Montag, D. Nasal Application of the Galantamine Pro-drug Memogain Slows Down Plaque Deposition and Ameliorates Behavior in 5X Familial Alzheimer’s Disease Mice. J. Alzheimers Dis. 46, 123–136 (2015).

6. Mucke, H. A. The case of galantamine: repurposing and late blooming of a cholinergic drug. Future Sci. OA 1, (2015).

7. Akram, M. N., Verpoorte, R. & Pomahačová, B. Effect of bulb age on alkaloid contents of narcissus pseudonarcissus bulbs. South Afr. J. Bot. 136, 182–189 (2021).

8. Marco-Contelles, J., do Carmo Carreiras, M., Rodríguez, C., Villarroya, M. & García, A. G. Synthesis and Pharmacology of Galantamine. Chem. Rev. 106, 116–133 (2006).

9. Fraser, M. D., Vallin, H. E., Davies, J. R. T., Rowlands, G. E. & Chang, X. Integrating Narcissus-derived galanthamine production into traditional upland farming systems. Sci. Rep. 11, 1389 (2021).

10. Effect of Fertilizers on Galanthamine and Metabolite Profiles in Narcissus Bulbs by 1H NMR | Journal of Agricultural and Food Chemistry. https://pubs-acs-org.ezproxy.lib.utexas.edu/doi/10.1021/jf104422m.

11. Thodey, K., Galanie, S. & Smolke, C. D. A microbial biomanufacturing platform for natural and semisynthetic opioids. Nat. Chem. Biol. 10, 837–844 (2014).

12. Payne, J. T., Valentic, T. R. & Smolke, C. D. Complete biosynthesis of the bisbenzylisoquinoline alkaloids guattegaumerine and berbamunine in yeast. Proc. Natl. Acad. Sci. 118, e2112520118 (2021).

13. Srinivasan, P. & Smolke, C. D. Biosynthesis of medicinal tropane alkaloids in yeast. Nature 585, 614–619 (2020).

14. Zhang, J. et al. A microbial supply chain for production of the anti-cancer drug vinblastine. Nature 609, 341–347 (2022).

15. Kilgore, M. B. & Kutchan, T. M. The Amaryllidaceae alkaloids: biosynthesis and methods for enzyme discovery. Phytochem. Rev. 15, 317–337 (2016).

16. Ehrenworth, A. M. & Peralta-Yahya, P. Accelerating the semisynthesis of alkaloid-based drugs through metabolic engineering. Nat. Chem. Biol. 13, 249–258 (2017).

17. d’Oelsnitz, S., et al. Using fungible biosensors to evolve improved alkaloid biosyntheses. Nat. Chem. Biol. 18, 981–989 (2022).

18. Schendzielorz, G. et al. Taking Control over Control: Use of Product Sensing in Single Cells to Remove Flux Control at Key Enzymes in Biosynthesis Pathways. ACS Synth. Biol. 3, 21–29 (2014).

19. Zhang, J. et al. Combining mechanistic and machine learning models for predictive engineering and optimization of tryptophan metabolism. Nat. Commun. 11, 4880 (2020).

20. Tang, S.-Y. et al. Screening for Enhanced Triacetic Acid Lactone Production by Recombinant Escherichia coli Expressing a Designed Triacetic Acid Lactone Reporter. J. Am. Chem. Soc. 135, 10099–10103 (2013).

21. Lu, H. et al. Machine learning-aided engineering of hydrolases for PET depolymerization. Nature 604, 662–667 (2022).

22. Hie, B. L. & Yang, K. K. Adaptive machine learning for protein engineering. Curr. Opin. Struct. Biol. 72, 145–152 (2022).

23. Greenhalgh, J. C., Fahlberg, S. A., Pfleger, B. F. & Romero, P. A. Machine learning-guided acyl-ACP reductase engineering for improved in vivo fatty alcohol production. Nat. Commun. 12, 5825 (2021).

24. Wu, Z., Kan, S. B. J., Lewis, R. D., Wittmann, B. J. & Arnold, F. H. Machine learning-assisted directed protein evolution with combinatorial libraries. Proc. Natl. Acad. Sci. U. S. A. 116, 8852–8858 (2019).

25. McNutt, A. T. et al. GNINA 1.0: molecular docking with deep learning. J. Cheminformatics 13, 43 (2021).

26. Jumper, J. et al. Highly accurate protein structure prediction with AlphaFold. Nature 596, 583–589 (2021).

27. Kilgore, M. B. et al. Cloning and Characterization of a Norbelladine 4′-O-Methyltransferase Involved in the Biosynthesis of the Alzheimer’s Drug Galanthamine in Narcissus sp. aff. pseudonarcissus. PLOS ONE 9, e103223 (2014).

28. Cravens, A., Payne, J. & Smolke, C. D. Synthetic biology strategies for microbial biosynthesis of plant natural products. Nat. Commun. 10, 2142 (2019).

29. Shroff, R., et al. Discovery of Novel Gain-of-Function Mutations Guided by Structure-Based Deep Learning. ACS Synth. Biol. 9, 2927–2935 (2020).

30. Paik, I. et al. Improved Bst DNA Polymerase Variants Derived via a Machine Learning Approach. Biochemistry 62, 410–418 (2023).

31. Kulikova, A. V., Diaz, D. J., Loy, J. M., Ellington, A. D. & Wilke, C. O. Learning the local landscape of protein structures with convolutional neural networks. J. Biol. Phys. 47, 435–454 (2021).

32. Stourac, J. et al. FireProtDB: database of manually curated protein stability data. Nucleic Acids Res. 49, D319–D324 (2021).

33. Newton, R. J., Hay, F. R. & Ellis, R. H. Temporal patterns of seed germination in early spring-flowering temperate woodland geophytes are modified by warming. Ann. Bot. 125, 1013–1023 (2020).

34. Zhang, Y., Ma, L., Su, P., Huang, L. & Gao, W. Cytochrome P450s in plant terpenoid biosynthesis: discovery, characterization and metabolic engineering. Crit. Rev. Biotechnol. 43, 1–21 (2023).

35. Noda, S. et al. Evaluation of Brachypodium distachyon L-Tyrosine Decarboxylase Using L-Tyrosine Over-Producing Saccharomyces cerevisiae. PLoS ONE 10, e0125488 (2015).

36. Curran, K. A., Leavitt, J. M., Karim, A. S. & Alper, H. S. Metabolic engineering of muconic acid production in Saccharomyces cerevisiae. Metab. Eng. 15, 55–66 (2013).

37. Abatemarco, J. et al. RNA-aptamers-in-droplets (RAPID) high-throughput screening for secretory phenotypes. Nat. Commun. 8, 332 (2017).

38. Kilgore, M. B., Holland, C. K., Jez, J. M. & Kutchan, T. M. Identification of a Noroxomaritidine Reductase with Amaryllidaceae Alkaloid Biosynthesis Related Activities*. J. Biol. Chem. 291, 16740–16752 (2016).

39. Singh, A. et al. Cloning and characterization of norbelladine synthase catalyzing the first committed reaction in Amaryllidaceae alkaloid biosynthesis. BMC Plant Biol. 18, 338 (2018).

40. Tousignant, L. et al. Transcriptome analysis of Leucojum aestivum and identification of genes involved in norbelladine biosynthesis. Planta 255, 30 (2022).

41. Kilgore, M. B., Augustin, M. M., May, G. D., Crow, J. A. & Kutchan, T. M. CYP96T1 of Narcissus sp. aff. pseudonarcissus Catalyzes Formation of the Para-Para’ C-C Phenol Couple in the Amaryllidaceae Alkaloids. Front. Plant Sci. 7, (2016).

