## Supporting Information for "Synthetic microbial sensing and biosynthesis of amaryllidaceae alkaloids"

**Supplementary Information:**

Supplementary Figures 1 - 11

Supplementary Tables 1-4

Supplementary Figures:

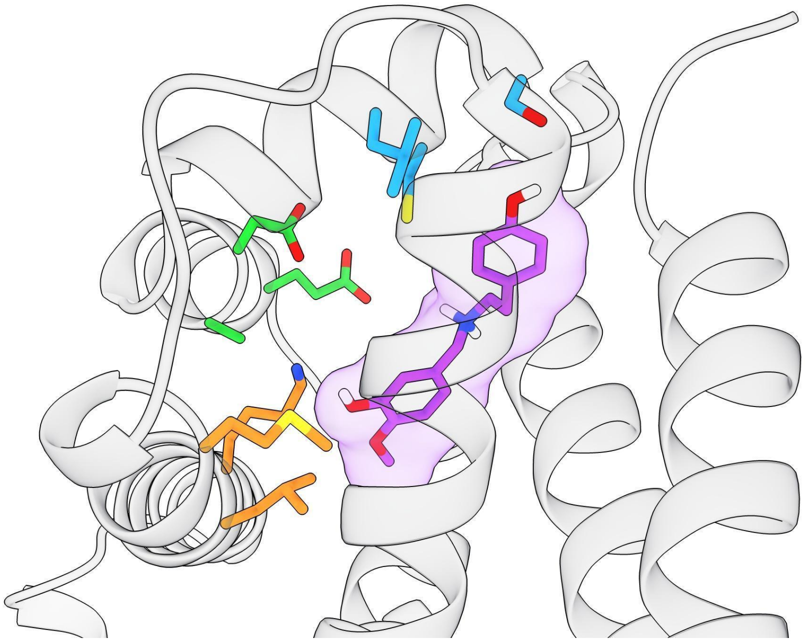

**Supplementary Figure 1**: Structural depiction of RamR library designs.
The structure of RamR (PDB: 3VVX) was docked with 4-OMe-norbelladine (purple) using the GNINA1.0 docking software. The side chains of residues targeted for site-saturation mutagenesis are color coded as follows. Orange: K63, L66, M71; Green: E120, A123, D124; Blue: L133, C134, S137.

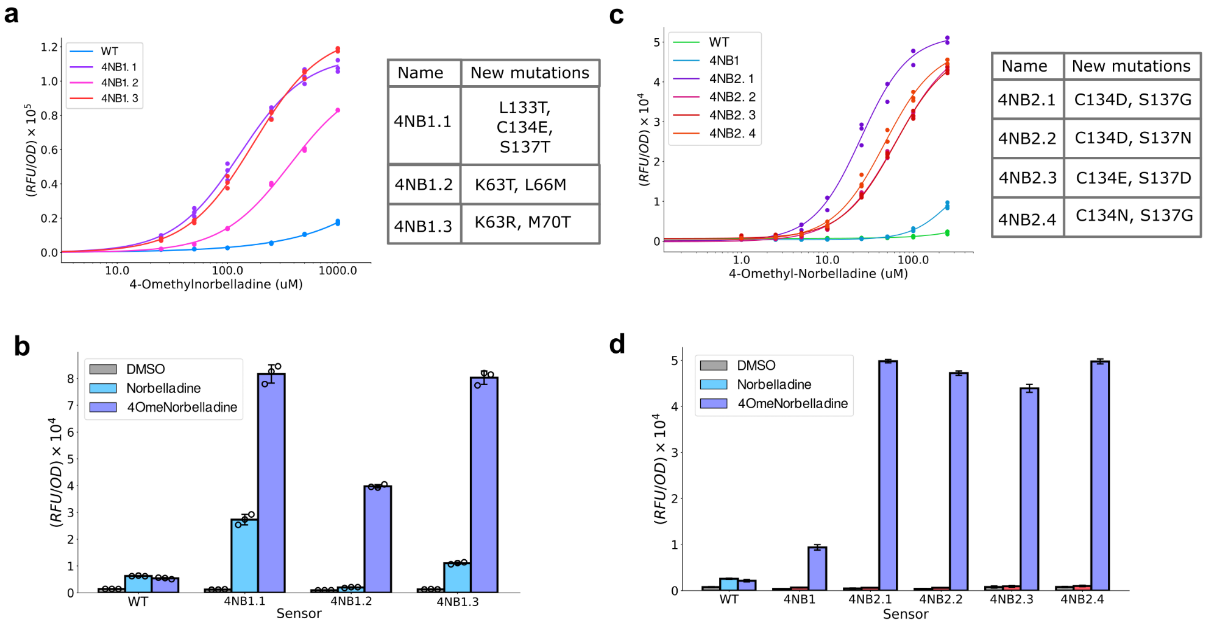


**Supplementary Figure 2:** The sensitivity and selectivity of RamR mutants evolved for 4-OMe-norbelladine. (a) Dose response measurements and genotype of generation one RamR sensors. (b) Selectivity

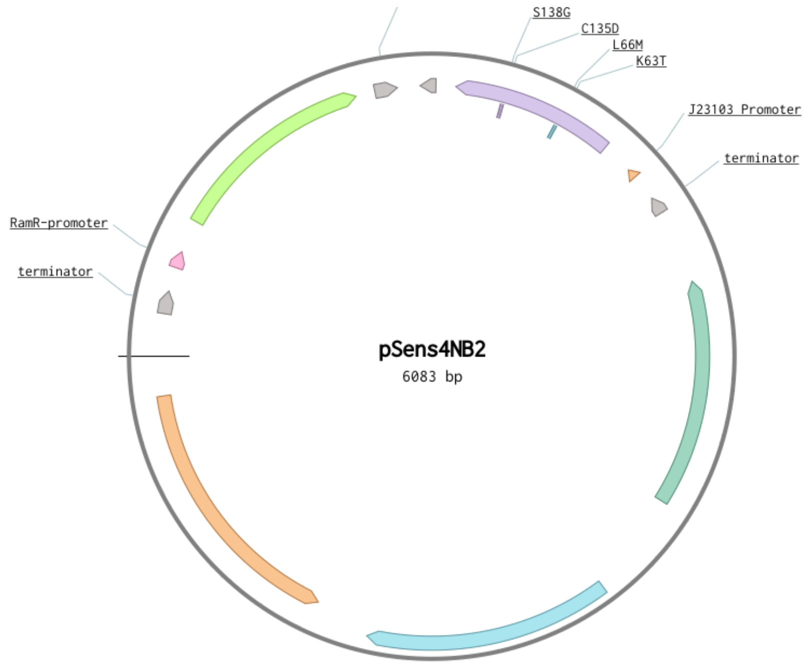


**Supplementary Figure 3:** Plasmid architecture for the one-plasmid 4-O'Methyl-norbelladine reporter system.

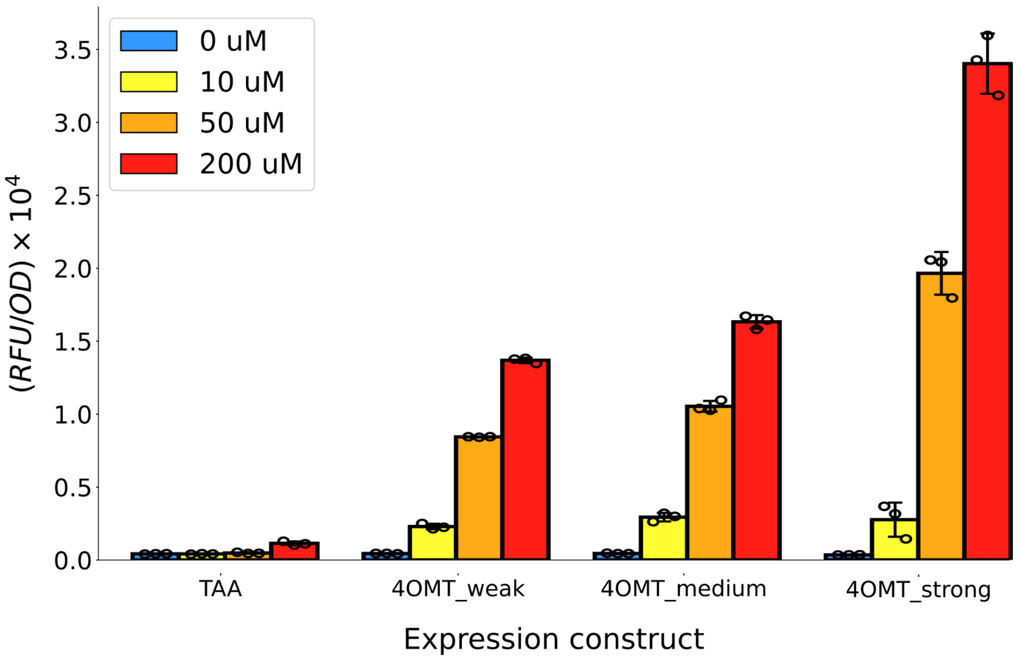


**Supplementary Figure 4:** Fluorescent response of 4NB2 with varying levels of Nb4OMT expression and precursor (norbelladine) supplementation. TAA represents an empty plasmid control in place of the Nb4OMT gene. Measurements were performed in biological triplicate and error bars represent the S.E.M. +/- the mean.

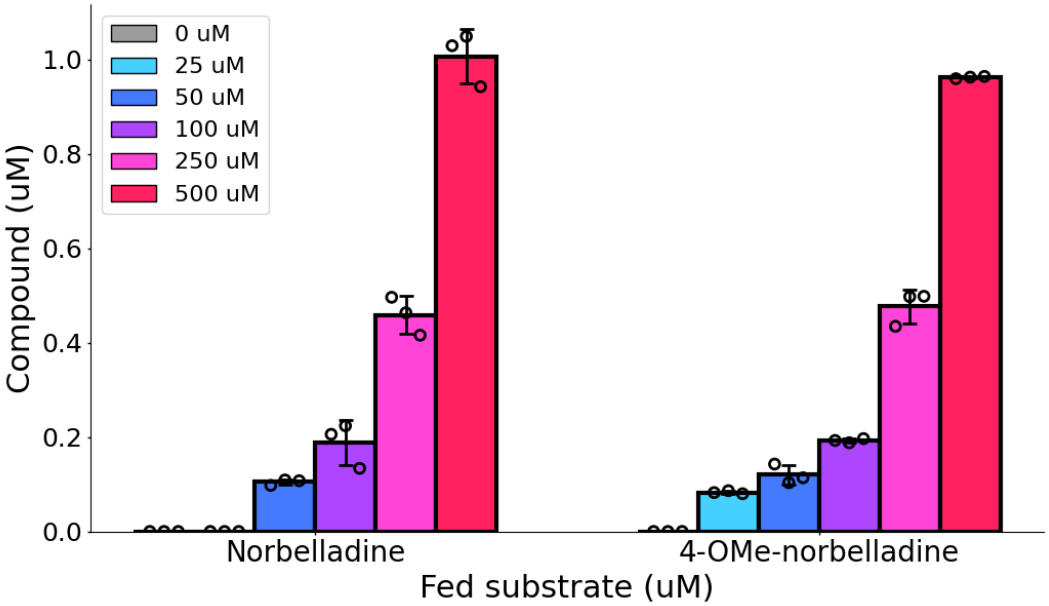


**Supplementary Figure 5:** Substrate and product of *in vivo* Nb4OMT reaction measured with HPLC. *E. coli* cells expressing the wild-type Nb4OMT enzyme were cultured with varying amounts of norbelladine for 18 hours and the concentrations of norbelladine and 4-OMe-norbelladine were subsequently measured. Measurements were performed in biological triplicate and error bars represent the S.E.M. +/- the mean.

**a**

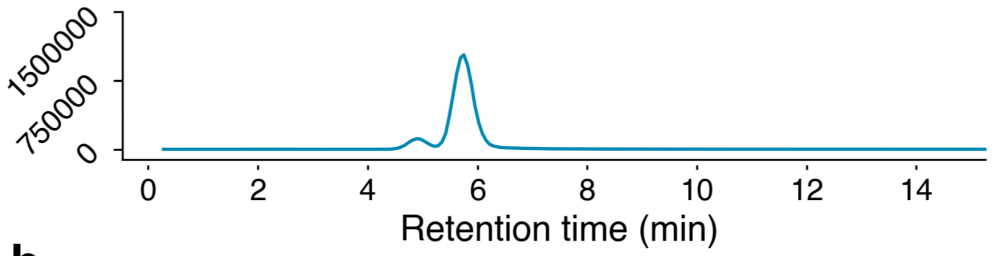


**b**

| Area Major Product | Area Minor Product | Major:Minor Ratio |
| --- | --- | --- |
| $1.23*10^8$ | $1.28*10^7$ | 9.59 |

**Supplementary Figure 6:** LC/MS analysis of the Nb4OMT-catalyzed *in vivo* reaction. *E. coli* cells expressing wild-type Nb4OMT were cultured for 24 hours with 500 uM of norbelladine and the culture supernatant was filtered and analyzed using LC/MS. (**a**) Ion-extracted chromatogram of the reaction product. The 274.1438 m/z ratio was used for extraction, since this m/z ratio is expected from all single methylated norbelladine products. (**b**) Statistics covering relative ion counts for the minor and major products.

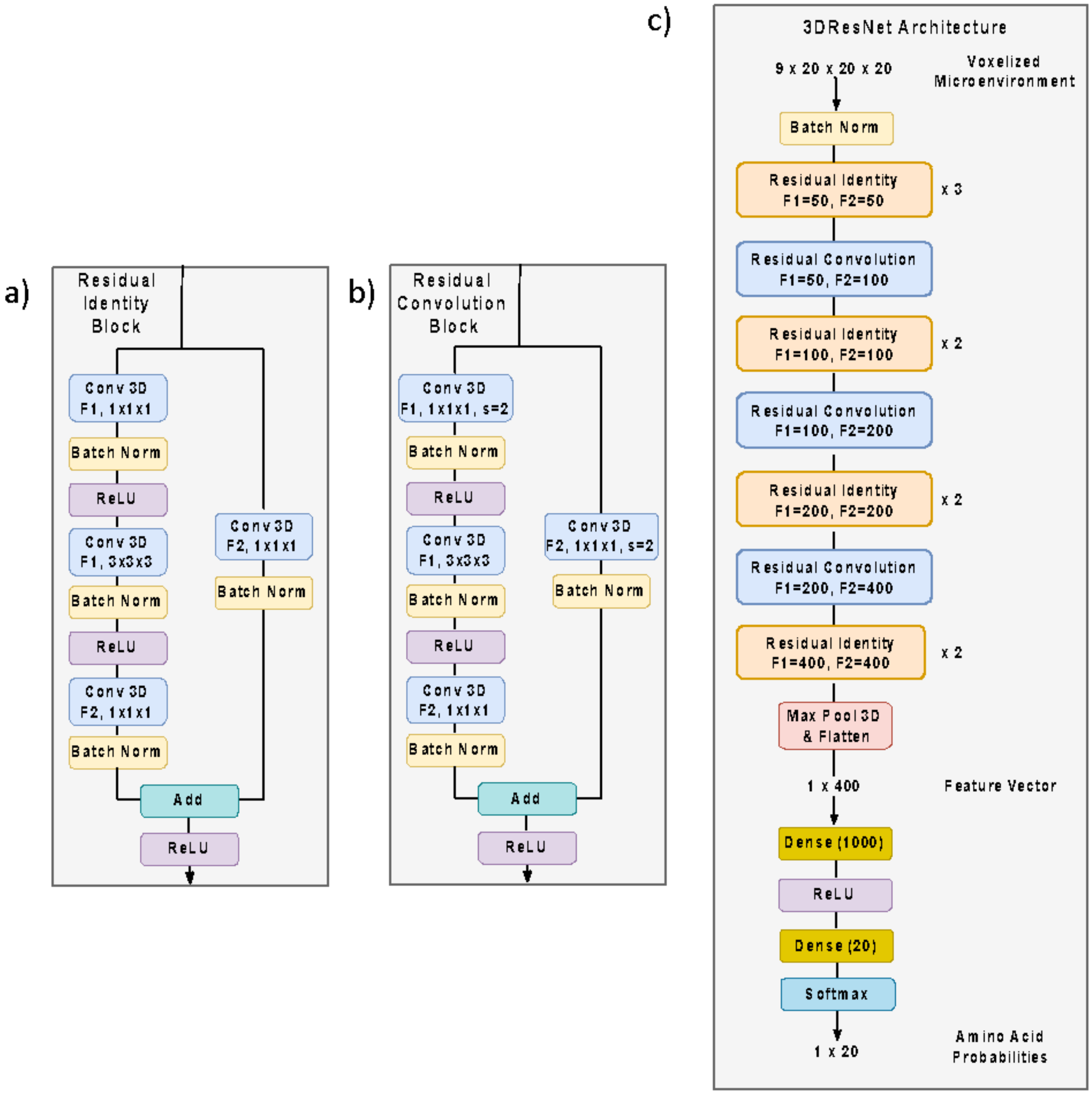


**Supplementary Figure 7.** Architecture details of 3DResNet: A) Residual Identity block B) Residual Convolutional block C) Full architecture of 3DResNet describing how a voxelized microenvironment (batch_size, 9 channels, 20x, 20y, 20z) is fed through the 3D residual feature extractor and converted into a 400-dimensional feature vector and then passed into a classifier to generate 20 amino acid probability distribution. Conv 3D: 3D Convolution layer, convolution kernel dimensions: 1x1x1 or 3x3x3, F1 and F2 are the number of feature maps generated by the convolution layer. S=2: stride of 2 used for that convolution layer else S=1 was used, ReLU: Rectified Linear Unit, Batch Norm: 3D Batch Normalization layer. Each convolution layer had L2 weight decay regularization set to 0.001. Batch normalization was instantiated with default hyperparameters.

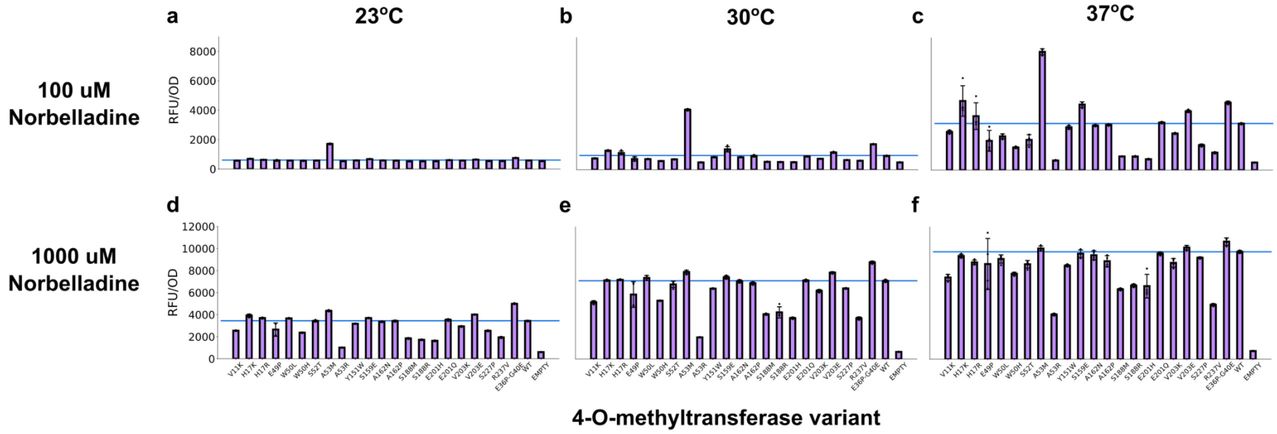


**Supplementary Figure 8**: Fluorescent response of cells containing single 4OMT variants across two precursor supplementation concentrations and three fermentation temperatures. In panels **a-c**, 100 uM of norbelladine was supplemented in the media, whereas for panels **d-f**, 1000 uM was supplemented. Measurements were performed in biological triplicate. Error bars represent the S.E.M. +/- the mean.

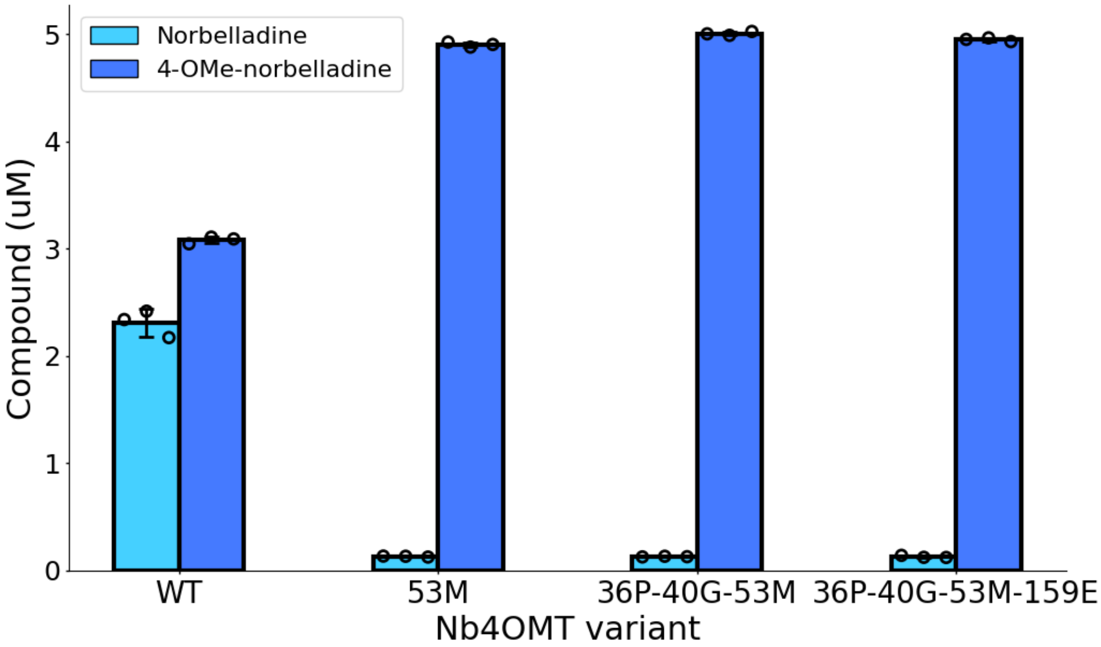


**Supplementary Figure 9:** HPLC-measured substrate and product concentrations resulting from *in vivo* reactions with Nb4OMT variants. Reactions were carried out within *E. coli* cells cultured for 24 hours at 37°C with 500 uM of norbelladine supplemented in the media. The resulting culture supernatant was filtered and compound concentrations were determined using HPLC. Mutations relative to the wild-type Nb4OMT sequence are labeled (for example, “53M”). Measurements were performed in biological triplicate. Error bars represent the S.E.M. +/- the mean.

**a**

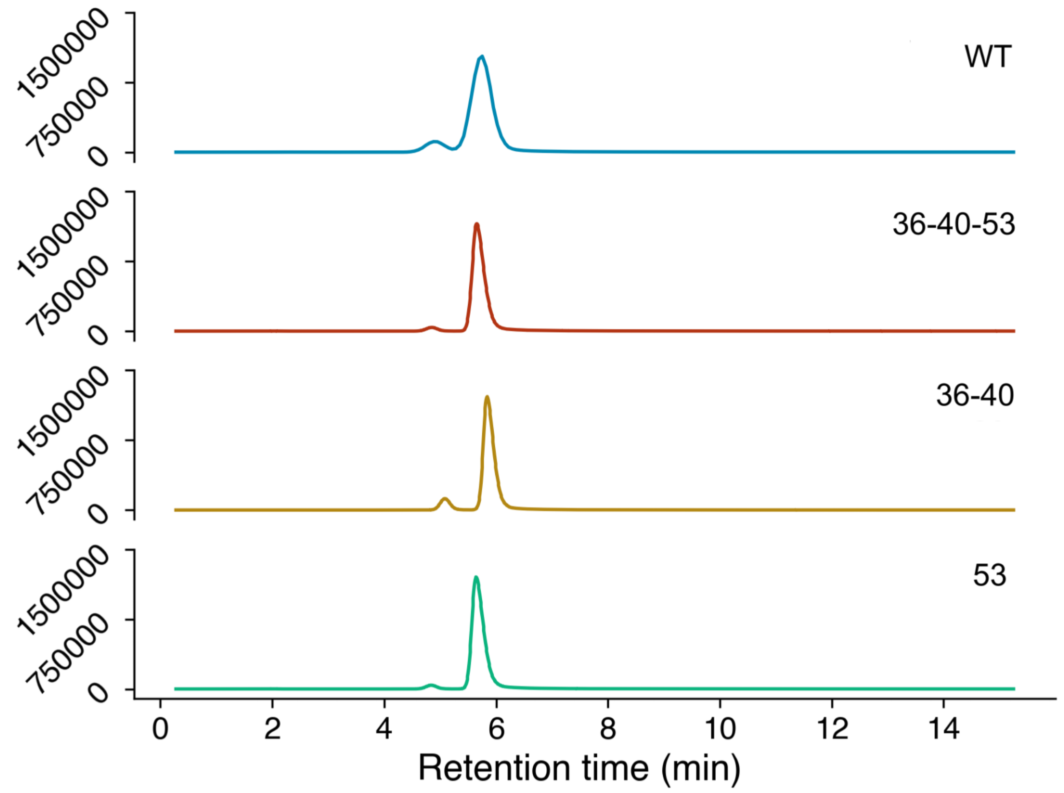


**b**

| Variant | Area Major Product | Area Minor Product | Major:Minor Ratio |
|---|---|---|---|
| WT | $1.28*10^8$ | $1.23*10^7$ | 9.59 |
| 36-40-53 | $1.85*10^7$ | $5.92*10^5$ | 31.32 |
| 36-40 | $1.71*10^7$ | $1.63*10^6$ | 10.50 |
| 53 | $1.90*10^7$ | $5.79*10^5$ | 32.75 |

**Supplementary Figure 10:** LC/MS analysis of the Nb4OMT variant-catalyzed *in vivo* reaction. *E. coli* cells expressing Nb4OMT variants were cultured for 24 hours with 500 uM of norbelladine and the culture supernatant was filtered and analyzed using LC/MS. (**a**) Ion-extracted chromatogram of the reaction product. The 274.1438 m/z ratio was used for extraction, since this m/z ratio is expected from all single methylated norbelladine products. (**b**) Statistics covering relative ion counts for the minor and major products. Variant name to mutation mapping is as follows, WT: the natural Nb4OMT sequence; 53: A53M; 36-40: E36P + G40E; 36-40-53: E36P + G40E + A53M.

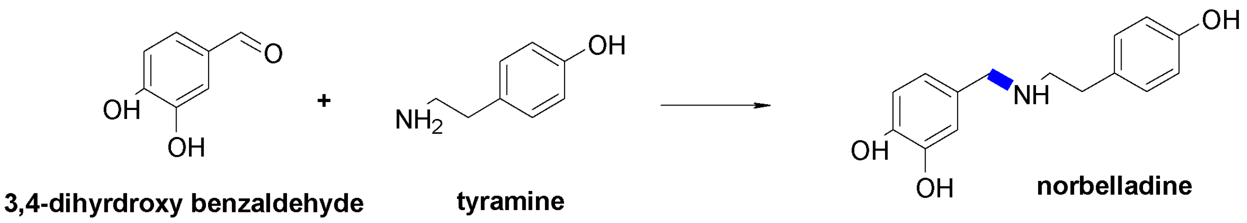


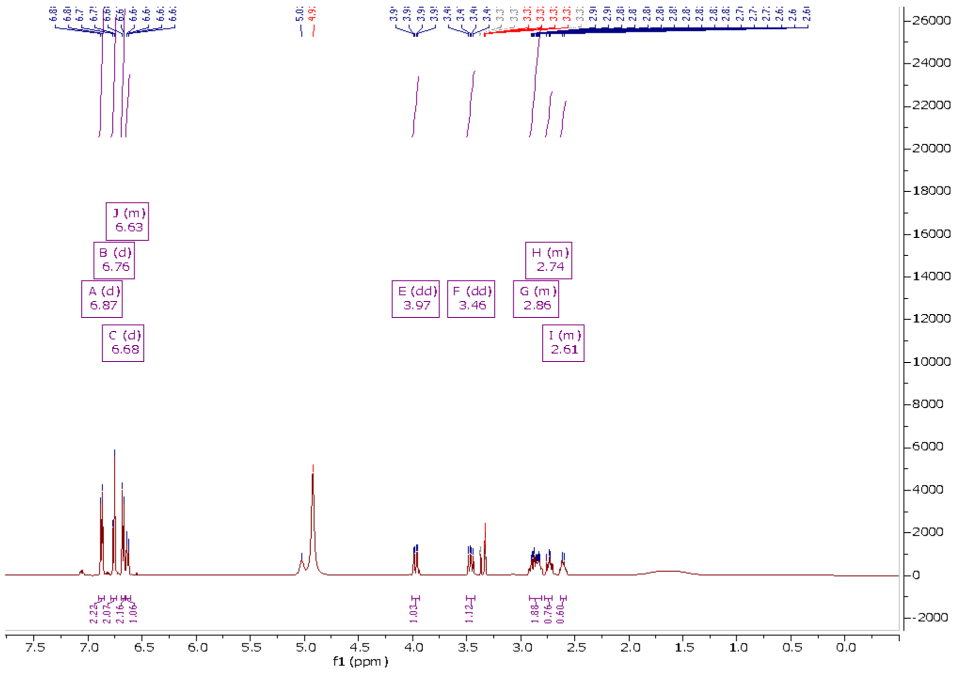


**Supplementary Figure 11:** Chemical synthesis and NMR validation of norbelladine. NMR spectra were taken on the 500 MHZ Bruker prodigy at University of Texas at Austin. NMR solvents (CD3OD). $^{1}$H NMR (500 MHz, MeOD) δ 6.87 (d, *J* = 8.2 Hz, 2H), 6.76 (m, *J* = 7.8 Hz, 2H), 6.67 (d, *J* = 8.2 Hz, 2H), 6.63 (d, *J* = 8.1 Hz, 1H), 3.97 (q, *J* = 5.6 Hz, 1H), 3.46 (q, *J* = 7.6 Hz, 1H), 2.87 (m, *J* = 4.6 Hz, 2H), 2.74 (m, *J* = 7.6 Hz, 1H), 2.61 (m, *J* = 3.5 Hz, 1H). Characterization was in accordance with previously reported literature.[1]

#### Supplementary Tables

| GNINA1.0 docking results | | | | |
|---|---|---|---|---|
| ligand | minimized Affinity (Kcal/mol) | Minimised RMSD (Angstrom) | CNN Score (probability) | CNN Affinity (pK) |
| S-adenosyl-Homocysteine | -7.918 | 0.557 | 0.835 | 5.851 |
| Norbelladine | -7.261 | 0.162 | 0.824 | 5.303 |

**Supplementary Table 1:** GNINA1.0 docking metrics for S-adenosyl-Homocysteine(SAH) and Norbelladine.

a)

| Interface Type | Percentage (%) |
|---|---|
| DNA | 0.04 |
| RNA | 0.72 |
| Ligand | 10.9 |
| Halogen | 1.17 |
| Protein | 23.98 |
| random core/surface | 63.19 |

b)

| Amino Acid | Percentage (%) |
|---|---|
| ALA | 7.84 |
| ARG | 5.51 |
| ASN | 4.39 |
| ASP | 6.01 |
| CYS | 1.4 |
| GLN | 3.72 |
| GLU | 6.68 |
| GLY | 7.18 |
| HIS | 2.71 |
| ILE | 5.6 |
| LEU | 9.08 |
| LYS | 5.6 |
| MET | 2.25 |
| PHE | 4.17 |
| PRO | 4.48 |
| SER | 5.89 |
| THR | 5.43 |
| TRP | 1.53 |
| TYR | 3.79 |
| VAL | 6.75 |

**Supplementary Table 2:** Dataset composition for training/testing 3DResNet models. Dataset consists of 22,759 protein sequences clustered at 50% sequence similarity from the PDB (as of November 2021). The sequences were sampled to generate a 2,569,256 microenvironment dataset, which was then split 90:10 to generate the training and test set, respectively. A) percentage of the dataset sampled at functional interfaces (within 5 angstroms of a non-protein atom or randomly from the core or surface of proteins. B) Amino acid composition of the dataset.

| Model | FP Wild Type Accuracy | Pearson ΔTm | Spearman ΔTm | Pearson ΔΔG | Spearman ΔΔG | ΔTm dataset size | ΔΔG dataset size |
|---|---|---|---|---|---|---|---|
| 3DCNN ensemble | 0.628 | 0.325 | 0.369 | -0.369 | -0.412 | 2719 | 4889 |
| 3DResNet ensemble | 0.597 | 0.367 | 0.425 | -0.408 | -0.457 | 2719 | 4889 |
| 3DResNet 1 | 0.626 | 0.332 | 0.383 | -0.381 | -0.429 | 2719 | 4889 |
| 3DResNet 2 | 0.656 | 0.405 | 0.426 | -0.421 | -0.452 | 2719 | 4889 |
| 3DResNet 3 | 0.591 | 0.378 | 0.435 | -0.423 | -0.474 | 2719 | 4889 |

**Supplementary Table 3:** Wildtype accuracy and Pearson and Spearman correlation metrics with ΔTm and ΔΔG experimental values for point mutations in FireProtDB[2] (as of February 2022). Correlations were calculated with the log odds predicted by the deep learning model for the mutated and wildtype amino acids: log $log\,(\frac{mutAA\_probability}{wtAA\_probability})$.

**>pSens4NB2**

TGGTTCGTTGTGATGGCGGTAGGAATGTAATCGTTAATCCGCAAATAACGTAAAAACCCGCTTCG
GCGGGTTTTTTTATGGGGGGAGTTTAGGGAAAGAGCATTTGTCATCCCGTTGAATATGGCTCGC
ATCTTATCGAGCATACTATCACGTCGGCGACCACTAGTCAGTTAACGCAAGGGCATGGGCTGTCG
ACCTTTGAAAAGTACCTTGACGGCGTATCTTTGCTTTCTATAATGAGTGCTTACTCACTCATACA
ATAGTCAGTCATAAGTCTGGGCTAAGCCCACTGATGAGTCGCTGAAATGCGACGAAACTTATGAC
CTCTACAAATAATTTTGTTTAACGTAAACCTCCGGGTTAATAAGGAGTAATTATGGCATCCAAGG
GCGAGGAGCTCTTTACTGGCGTAGTACCAATTCTCGTAGAGCTCGATGGCGATGTAAATGGCCAT
AAGTTTTCCGTACGCGGCGAGGGCGAGGGCGATGCAACTAACGGCAAGCTCACTCTCAAGTTTA
TTTGTACTACTGGCAAGCTCCCAGTACCATGGCCAACTCTCGTAACTACTCTGACCTATGGCGTAC
AATGTTTTTCCCGCTATCCAGATCACATGAAGCAACATGATTTTTTTAAGTCCGCAATGCCAGAG
GGCTATGTACAAGAGCGCACTATTAGCTTTAAGGATGATGGCACCTATAAGACTCGCGCAGAGGT
AAAGTTTGAGGGCGATACTCTCGTAAATCGCATTGAGCTCAAGGGCATTGATTTTAAGGAGGAT
GGCAATATTCTCGGCCATAAGCTGGAGTATAATTTCAATTCCCATAATGTATATATTACCGCAGAT
AAGCAAAAGAATGGCATTAAGGCGAATTTTAAGATTCGCCATAATGTGGAGGATGGCTCCGTAC
AACTCGCAGATCATTATCAACAAAATACTCCAATTGGCGATGGCCCAGTACTCCTCCCAGATAAT
CATTATCTCTCCACTCAATCCGTGCTCTCCAAAGATCCAAATGAGAAGCGCGATCACATGGTACT
CCTGGAGTTTGTAACTGCAGCAGGCATTACTCATGGCATGGATGAGCTCTATAAGCTCGAGCACC
ACCACCACCACCACTGATAATCCAAACCTGTTATATGTTAGCTGAGACTAGTTGGAAGTGTGGCT
GTCCTCAAGCGTTTTAGTTCGTCGGTCAGTTTCACCTGATTTACGTAAAAACCCGCTTCGGCGGG
TTTTTGCTTTTGGAGGGGCAGAAAGATGAATGACTGTCGGCCATTCGATGGTGTCGGGTAGCAT
AACCCCTTGTGATACCTTTGCCATGTTTCAGAAACAACTCTGGCGCATCGGGCTTGGACCAAAA
CGAAAAAAGGCCGCTTTCGCGGCCTCTTTTCTGGAATTTGGTACCGAGCTCACAGCGGCGATAG
TCAGATAGCTAGACCGTATGTTACCGAGCCTGCACTCCTCGAATTCTTAGGATTATTACTGCTCT
TCGCGCGTAAGTGCGCGCCACATAGCCTCGAAGCCCAACGCAATGTACTCACCAGCGCGAGCCGG
GTCGCGCGCAGCGAAATCCATAGTCGTCTCAGCAAGCGCCAAGAACAACCCGTCGCCGAAGGCGC
GGTACTCGTCGGACATAAACACCATAAGAACGCCACGGTGGTCGAGGTCGCGTAACTCCGGGAAC
ATATCATCCGCGCGTTGTTCGGTTTCCTTCGTCAACTTTTCAGAAACCGCCAACTGACGAATGGC
ACGATGGCGAGCTGGGTGGTTCAATCCCCAGCTAATATAACTGTTCCAGATAAAACGGGTCATCA
TCTTAGCGTCAGTAATAGAACGATCCAATTCCATGATCATTGATTGGCACATGTCCTGGGTCAAA
TGTAAGTAAAGGGTGTTGATCAACTCATCTTTCGTTGCGAAATAGCGGAACAACGTCCCTTCCGC
AACTCCCGCATTGCGTGCAATTACAGCGGTACTAGCGGCAATGCCTGATTGCGCGATGGCTTGAG
TTGCCGCTTCAAGCAATGCCTGCTTTTTGTCCTCAGACTTTGGGCGAGCAACCATATACTAACCT
CCTTCTGATACGTGGTTCCGTTAAACAAAATTATTTGTAGAGGCCCCATTTCGTCCTTTTGGACT
CATCAGGGGTGGTACACACCACCCTATGGGGCTCGTAATTGCTAGCATAATCCCTAGGACTGAGC
TAGCTATCAGGGTACTTTTCAAAGGTCGACAGCCCATGCCCTTGCGTTCGGCAGGTGTACAATGA
TACGAGGTAATGAAGATGAAGTCCATACAATCGATAGATTGGGACCAAAACGAAAAAAGGGGAG
CGGTTTCCCGCTCCCCTCTTTTCTGGAATTTGGTACCGAGTCGCACCTGATTGCCCGACATTATC
GCACGGTGTCTCATCTCTGATAACGCATATTGTCGTTAGAACTCGGCGCGGCCGCTCACACTGCT
TCCGGTAGTCAATAAACCGGTAAACCAGCAATAGACATAAGCGGCTATTTAACGACCCTGCCCTG
AACCGACGACCGGGTCGAATTTGCTTTCGAATTTCTGCCATTCATCCGCTTATTATCACTTATTC
AGGCGTAGCAACCAGGCGTTTAAGGGCACCAATAACTGCCTTAAAAAAATTAGAAAAACTCATC
GAGCATCAAATGAAACTGCAATTTATTCATATCAGGATTATCAATACCATATTTTTGAAAAAGCC
GTTTCTGTAATGAAGGAGAAAACTCACCGAGGCAGTTCCATAGGATGGCAAGATCCTGGTATCG
GTCTGCGATTCCGACTCGTCCAACATCAATACAACCTATTAATTTCCCCTCGTCAAAAATAAGGT
TATCAAGTGAGAAATCACCATGAGTGACGACTGAATCCGGTGAGAATGGCAAAAGTTTATGCAT
TTCTTTCCAGACTTGTTCAACAGGCCAGCCATTACGCTCGTCATCAAAATCACTCGCATCAACCA
AACCGTTATTCATTCGTGATTGCGCCTGAGCGAGACGAAATACGCGGTCGCTGTTAAAAGGACA

ATTACAAACAGGAATCGAATGCAACCGGCGCAGGAACACTGCCAGCGCATCAACAATATTTTCA
CCTGAATCAGGATATTCTTCTAATACCTGGAATGCTGTTTTCCCGGGGATCGCAGTGGTGAGTAA
CCATGCATCATCAGGAGTACGGATAAAATGCTTGATGGTCGGAAGAGGCATAAATTCCGTCAGCC
AGTTTAGTCTGACCATCTCATCTGTAACATCATTGGCAACGCTACCTTTGCCATGTTTCAGAAAC
AACTCTGGCGCATCGGGCTTCCCATACAATCGATAGATTGTCGCACCTGATTGCCCGACATTATC
GCGAGCCCATTTATACCCATATAAATCAGCATCCATGTTGGAATTTAATCGCGGCCTAGAGCAAG
ACGTTTCCCGTTGAATATGGCTCATTTTAGCTTCCTTAGCTCCTGAAAATCTCGATAACTCAAAA
AATACGCCCGGTAGTGATCTTATTTCATTATGGTGAAAGTTGGAACCTCTTACGTGCCGATCACG
TCTCATTTTCGCCAAAGTTGGCCAGGGCTTCCCGGTATCAACAGGGACACCAGGATTTATTTATN
NTGCGAAGTGATCTTCCGTCACAGGTATTTATTCGGCGCAAAGTGCGTCGGGTGATGCTGCCAA
CTTACTGATTTAGTGTATGATGGTGTTTTTGAGGTGCTCCAGTGGCTTCTGTTTCTATCAGCTGT
CCCTCCTGTTCAGCTACTGACGGGGTGGTGCGTAACGGCAAAAGCACCGCCGGACATCAGCGCTA
GCGGAGTGTATACTGGCTTACTATGTTGGCACTGATGAGGGTGTCAGTGAAGTGCTTCATGTGGC
AGGAGAAAAAAGGCTGCACCGGTGCGTCAGCAGAATATGTGATACAGGATATATTCCGCTTCCTC
GCTCACTGACTCGCTACGCTCGGTCGTTCGACTGCGGCGAGCGGAAATGGCTTACGAACGGGGC
GGAGATTTCCTGGAAGATGCCAGGAAGATACTTAACAGGGAAGTGAGAGGGCCGCGGCAAAGCC
GTTTTTCCATAGGCTCCGCCCCCCTGACAAGCATCACGAAATCTGACGCTCAAATCAGTGGTGGC
GAAACCCGACAGGACTATAAAGATACCAGGCGTTTCCCCCTGGCGGCTCCCTCGTGCGCTCTCCT
GTTCCTGCCTTTCGGTTTACCGGTGTCATTCCGCTGTTATGGCCGCGTTTGTCTCATTCCACGCC
TGACACTCAGTTCCGGGTAGGCAGTTCGCTCCAAGCTGGACTGTATGCACGAACCCCCCGTTCAG
TCCGACCGCTGCGCCTTATCCGGTAACTATCGTCTTGAGTCCAACCCGGAAAGACATGCAAAAGC
ACCACTGGCAGCAGCCACTGGTAATTGATTTAGAGGAGTTAGTCTTGAAGTCATGCGCCGGTTAA
GGCTAAACTGAAAGGACAAGTTTTGGTGACTGCGCTCCTCCAAGCCAGTTACCTCGGTTCAAAG
AGTTGGTAGCTCAGAGAACCTTCGAAAAACCGCCCTGCAAGGCGGTTTTTTCGTTTTCAGAGCA
AGAGATTACGCGCAGACCAAAACGATCTCAAGAAGATCATCTTATTAATCAGATAAAATATTTCT
AGATTTCAGTGCAATTTATCTCTTCAAATGTAGCACCTGAAGTCAGCCCCATACGATATAAGTTG
TAATTCTCATGTTAGTCATGCCCCGCGCCCACCGGAAGGAGCTGACTGGGTTGAAGGCTCTCAAG
GGCATCGGTCGAGATCCCGGTGCCTAATGAGTGAGCTAACTTACATTAATTGCGTTGCGCTCACT
GCCCGCTTTCCAGTCGGGAAACCTGTCGTGCCAGCTGCATTAATGAATCGGCCAACGCGCGGGGA
GAGGCGGTTTGCGTATTGGGCGCCAGGGTGGTTTTTCTTTTCACCAGTGAGACGGGCAACAGCT
GATTGCCCTTCACCGCCTGGCCCTGAGAGAGTTGCAGCAAGCGGTCCACGCTGGTTTGCCCCAGC
AGGCGAAAATCCTGTTTGATGGTGGTTAACGGCGGGATATAACATGAGCTATCTTCGGTATCGTC
GTATCCCACTACCGAGATGTCCGCACCAACGCGCAGCCCGGACTCGGTAATGGCGCGCATTGCGC
CCAGCGCCATCTGATCGTTGGCAACCAGCATCGCAGTGGGAACGATGCCCTCATTCAGCATTTGC
ATGGTTTGTTGAAAACCGGACATGGCACTCCAGTCGCCTTCCCGTTCCGCTATCGGCTGAATTTG
ATTGCGAGTGAGATATTTATGCCAGCCAGCCAGACGCAGACGCGCCGAGACAGAACTTAATGGGC
CCGCTAACAGCGCGATTTGCTGGTGACCCAATGCGACCAGATGCTCCACGCCCAGTCGCGTACCA
TCTTCATGGGAGAAAATAATACTGTTGATGGGTGTCTGGTCAGAGACATCAAGAAATAACGCCG
GAACATTAGTGCAGGCAGCTTCCACAGCAATGGCATCCTGGTCATCCAGCGGATAGTTAATGATC
AGCCCACTGACGCGTTGCGCGAGAAGATTGTGCACCGCCGCTTTACAGGCTTCGACGCCGCTTCG
TTCTACCATCGACACCACCACGCTGGCACCCAGTTGATCGGCGCGAGATTTAATCGCCGCGACAA
TTTGCGACGGCGCGTGCAGGGCCAGACTGGAGGTGGCAACGCCAATCAGCAACGACTGTTTGCC
CGCCAGTTGTTGTGCCACGCGGTTGGGAATGTAATTCAGCTCCGCCATCGCCGCTTCCACTTTTT
CCCGCGTTTTCGCAGAAACGTGGCTGGCCTGGTTCACCACGCGGGAAACGGTCTGATAAGAGAC
ACCGGCATACTCTGCGACATCGTATAACGTTACTGGTTTCACATTCACCACCCTGAATTGACTCT
CTTCCGGGCGCTATCATGCCATACCGCGAAAGGTTTTGCGCCATTCGATGGTGTCCGGGATCTCG
ACGCTCTCCCTTATGCGACGCGGCCGCGGCATCAGAGCAGATTGTACTGTGTCCTCAA

>Nb4OMT-A53M
ATCCCCCTTACACGGAGGCATCAGTGACCAAACAGGAAAAAACCGCCCTTAACATGGCCCGCTTT
ATCAGAAGCCAGACATTAACGCTTCTGGAGAAACTCAACGAGCTGGACGCGGATGAACAGGCAG
ACATCTGTGAATCGCTTCACGACCACGCTGATGAGCTTTACCGCAGCTGCCTCGCGCGTTTCGGT
GATGACGGTGAAAACCTCTGACACATGCAGCTCCCGCAGACGGTCACAGCTTGTCTGTAAGCGG
ATGCCGGGAGCAGACAAGCCCGTCAGGGCGCGTCAGCGGGTGTTGGCGGGTGTCGGGGCGCAGC
CATGACCCAGTCACGTAGCGATAGCGGAGTGTATACTGGCTTAACTATGCGGCATCAGAGCAGAT
TGTACTGAGAGTGCACCGGTGTGAAATACCGCACAGATGCGTAAGGAGAAAATACCGCATCAGGC
GCTCTTCCGCTTCCTCGCTCACTGACTCGCTGCGCTCGGTCGTTCGGCTGCGGCGAGCGGTATCA
GCTCACTCAAAGGCGGTAATACGGTTATCCACAGAATCAGGGGATAACGCAGGAAAGAACATGT
GAGCAAAAGGCCAGCAAAAGGCCAGGAACCGTAAAAAGGCCGCGTTGCTGGCGTTTTTCCATAG
GCTCCGCCCCCCTGACGAGCATCACAAAAATCGACGCTCAAGTCAGAGGTGGCGAAACCCGACA
GGACTATAAAGATACCAGGCGTTTCCCCCTGGAAGCTCCCTCGTGCGCTCTCCTGTTCCGACCCT
GCCGCTTACCGGATACCTGTCCGCCTTTCTCCCTTCGGGAAGCGTGGCGCTTTCTCATAGCTCAC
GCTGTAGGTATCTCAGTTCGGTGTAGGTCGTTCGCTCCAAGCTGGGCTGTGTGCACGAACCCCCC
GTTCAGCCCGACCGCTGCGCCTTATCCGGTAACTATCGTCTTGAGTCCAACCCGGTAAGACACGA
CTTATCGCCACTGGCAGCAGCCACTGGTAACAGGATTAGCAGAGCGAGGTATGTAGGCGGTGCTA
CAGAGTTCTTGAAGTGGTGGCCTAACTACGGCTACACTAGAAGGACAGTATTTGGTATCTGCGCT
CTGCTGAAGCCAGTTACCTTCGGAAAAAGAGTTGGTAGCTCTTGATCCGGCAAACAAACCACCG
CTGGTAGCGGTGGTTTTTTTGTTTGCAAGCAGCAGATTACGCGCAGAAAAAAAGGATCTCAAGA
AGATCCTTTGATCTTTTCTACGGGGTCTGACGCTCAGTGGAACGAAAACTCACGTTAAGGCCCTC
TCCAAGACCGAGCCATCAACAAAGCGTCTCGCTGAGGTTTCATGGAGCCTCTGGTTCATCTCCGG
CAATTAAAAAAGCGGCTAACCACGCCGCTTTTTTTACGTCTGCAGGAACGGGCTGTCGACCTTT
GAAAAGTTCGTTTACCGCTAGCTCAGTCCTAGGTACAATTACAGCCATCGTACGAGCCCTGGCTG
AGCACAGCTGTCACCGGATGTGCTTTCCGGTCTGATGAGTCCGTGAGGACGAAACAGCCTCTACA
AATAATTTTGTTTAACTAGTGAACCACGAGGCCTACATATGGGTGCTTCAATTGATGACTACTC
CTTGGTACATAAAAACATCTTGCATTCCGAAGATCTGCTGAAATACATTCTTGAAACCAGTGCAT
ATCCTCGCGAACACGAACAATTAAAGGGTCTGCGTGAGGTTACTGAAAAGCACGAATGGTCATC
CatgTTAGTACCGGCAGACGAAGGTTTATTCCTGTCAATGTTACTGAAGTTGATGAATGCAAAAC
GTACTATCGAAATCGGCGTGTACACGGGATACAGCCTGTTAACAACGGCATTGGCTTTACCGGAG
GATGGTAAAATTACGGCGATCGATGTAAATAAGAGTTATTACGAAATCGGATTGCCCTTTATTCA
GAAGGCGGGCGTGGAACACAAGATCAACTTTATTGAGTCGGAAGCGCTTCCCGTGCTGGATCAA
ATGTTAGAGGAGATGAAGGAGGAGGATTTATACGACTACGCTTTTGTGGATGCTGATAAAAGCA
ATTATGCCAATTACCATGAGCGTCTTGTAAAACTTGTACGTATCGGTGGTGCCATCCTGTACGAC
AATACACTGTGGTATGGTTCTGTTGCGTACCCGGAATACCCCGGTCTGCATCCAGAGGAAGAAGT
CGCGCGTCTGAGCTTTCGTAACTTGAATACCTTTTTAGCAGCAGATCCTCGCGTAGAAATTAGTC
AAGTCTCAATTGGTGATGGCGTGACCATTTGTCGCCGCTTGTATTAATAATCCTATCGCCACTTT
CAGCCAAAAAACTTAAGACCGCCGGTCTTGTCCACTACCTTGCAGTAATGCGGTGGACAGGATCG
GCGGTTTTCTTTTCTCTTCTCAACACCCTTCGCGTCAACACTTTTCCGCCAAGGAGACGGTTGGT
CAGGTTTTCGGGAGGTGTGGCTGGAAGTTCCTATACTTTCTAGAGAATAGGAACTTCTTTCTAA
ATACATTCAAATATGTATCCGCTCATGAGACAATAACCCTGATAAATGCTTCAATAATATTGAAA
AAGGAAGAGTATGAGTATTCAACATTTCCGTGTCGCCCTTATTCCCTTTTTTGCGGCATTTTGCC
TTCCTGTTTTTGCTCACCCAGAAACGCTGGTGAAAGTAAAAGATGCTGAAGATCAGTTGGGTGC
ACGAGTGGGTTACATCGAACTGGATCTCAACAGCGGTAAGATCCTTGAGAGTTTTCGCCCCGAA
GAACGTTTTCCAATGATGAGCACTTTTAAAGTTCTGCTATGTGGCGCGGTATTATCCCGTGTTGA
CGCCGGGCAAGAGCAACTCGGTCGCCGCATACACTATTCTCAGAATGACTTGGTTGAGTACTCAC
CAGTCACAGAAAAGCATCTTACGGATGGCATGACAGTAAGAGAATTATGCAGTGCTGCCATAACC
ATGAGTGATAACACTGCGGCCAACTTACTTCTGACAACGATCGGAGGACCGAAGGAGCTAACCGC

| |
|---|
| TTTTTTGCACAACATGGGGGATCATGTAACTCGCCTTGATCGTTGGGAACCGGAGCTGAATGAA<br>GCCATACCAAACGACGAGCGTGACACCACGATGCCTGCAGCAATGGCAACAACGTTGCGCAAAC<br>TATTAACTGGCGAACTACTTACTCTAGCTTCCCGGCAACAATTAATAGACTGGATGGAGGCGGAT<br>AAAGTTGCAGGACCACTTCTGCGCTCGGCCCTTCCGGCTGGCTGGTTTATTGCTGATAAATCTGG<br>AGCCGGTGAGCGTGGATCGCGCGGTATCATTGCAGCACTGGGGCCAGATGGTAAGCCCTCCCGTA<br>TCGTAGTTATCTACACGACGGGGAGTCAGGCAACTATGGATGAACGAAATAGACAGATCGCTGAG<br>ATAGGTGCCTCACTGATTAAGCATTGGTAAGTTGTGATGGCGGTAGGAATGTAATCGTTAATCCG<br>CAAATAACGTAAAAACCCGCTTCGGCGGGTTTTTTTATGGGGGGAGTTTAGGGAAAGAGCATTT<br>GTCATCCCGTTGAATATGGCTCCCTTAACGTGAGGAAGTTCCTATACTTTCTAGAGAATAGGAAC<br>TTCTACAGATGGACTTGGGTTGGCGGTTTCAGGAGTCTGCAAAACGTCTGCGACCTGAGCAACA<br>ACATGAATGGTCATCGGTTTCCGTGTTTCGTAAAGTCTGGAAACGCGGAAGTCAGCGCCCTGCA<br>CCATTATGTTCCGGATCTGCATCGCAGGATGCTGCTGGCTACCCTGTGGAACACCTACATCTGTA<br>TTAACGAAGCGCTGGCATTGACCCTGAGTGATTTTTCTCTGGTCCCGCCGCATCCATACCGCCAG<br>TTGTTTACCCTCACAACGTTCCAGTAACCGGGCATGTTCATCATCAGTAACCCGTATCGTGAGCA<br>TCCTCTCTCGTTTCATCGGTATCATTACCCCCATGAACAGAA |
| **Sequence annotation color code map:**<br>LacI<br>p15A origin<br>Kanamycin resistance<br>Terminator<br>RamR-4NB2<br>GFP<br>RamR promoter<br>pBR322 origin<br>Ampicillin Resistance |

**Supplementary Table 4:** Sequences with color-coded annotations of plasmids used in this study.

### References

1. Park, J. B. Synthesis and characterization of norbelladine, a precursor of Amaryllidaceae alkaloid, as an anti-inflammatory/anti-COX compound. *Bioorganic & Medicinal Chemistry Letters* **24**, 5381–5384 (2014).

2. Stourac, J., Dubrava, J., Musil, M., Horackova, J., Damborsky, J., Mazurenko, S., Bednar, D., 2020: FireProt$^{DB}$: Database of Manually Curated Protein Stability Data. *Nucleic Acids Research* 49: D319-D324
